# Effective Inclusion of Electronic Polarization Improves the Description of Electrostatic Interactions: The prosECCo75 Biomolecular Force Field

**DOI:** 10.1101/2024.05.31.596781

**Authors:** Ricky Nencini, Carmelo Tempra, Denys Biriukov, Miguel Riopedre-Fernandez, Victor Cruces Chamorro, Jakub Polák, Philip E. Mason, Daniel Ondo, Jan Heyda, O. H. Samuli Ollila, Pavel Jungwirth, Matti Javanainen, Hector Martinez-Seara

## Abstract

`prosECCo75` is an optimized force field effectively incorporating electronic polarization via charge scaling. It aims to enhance the accuracy of nominally nonpolarizable molecular dynamics (MD) simulations for interactions in biologically relevant systems involving water, ions, proteins, lipids, and saccharides. Recognizing the inherent limitations of nonpolarizable force fields in precisely modeling electrostatic interactions essential for various biological processes, we mitigate these shortcomings by accounting for electronic polarizability in a physical rigorous mean-field way that does not add to computational costs. With this scaling of (both integer and partial) charges within the CHARMM36 framework, `prosECCo75` addresses overbinding artifacts. This improves agreement with experimental ion binding data across a broad spectrum of systems — lipid membranes, proteins (including peptides and amino acids), and saccharides — without compromising their biomolecular structures. `prosECCo75` thus emerges as a computationally efficient tool providing enhanced accuracy and broader applicability in simulating the complex interplay of interactions between ions and biomolecules, pivotal for improving our understanding of many biological processes.

## 1 Introduction

Understanding the complexity of cellular structures at the molecular scale is instrumental in better comprehending biological processes and designing more potent and specific drugs. Although temporal and spatial resolutions of experimental techniques are steadily improving, computational methods such as molecular dynamics (MD) simulations still constitute the most detailed “atomistic microscope”.^1^ MD simulations can track the movement of individual atoms in systems ranging from simple aqueous solutions all the way to realistic cell membranes or protein complexes, which are presently central study targets in biosciences. The ability of MD simulations to capture the interplay between water, ions, proteins, lipids, and polysaccharides at a resolution hardly accessible *in vitro* — let alone *in vivo* — thus offers a viable alternative performing computer experiments *in silico* instead. One of the main challenges for simulations is to describe with sufficient accuracy interactions involving biomolecules, water, ions, and other solutes. Biomolecular force fields have witnessed a steady improvement in their accuracy throughout the years, thanks to refinement efforts by multiple research groups.^2–6^ Still, the increasing complexity of systems that can be simulated nowadays calls for a careful balancing of the interactions among an ever-increasing number of types of molecules.

Electrostatic interactions play a crucial role in various biological processes, such as for example intercellular signaling mediated by Ca^2+^ ions,^7^ stabilization of protein structures through salt bridges,^8^ enzyme activity relying on poly-coordinated ions,^9^ or the adsorption of peripheral proteins to charged membranes.^10^ For membrane-involving processes in particular, the importance of electrostatics is highlighted in the vicinity of the intracellular leaflet, where anionic lipids and ions are involved in the signaling by charged proteins. ^10^

One of the key aspects modulating the electrostatic interactions between charged molecular groups is electronic polarizability.^11^ The lack of its description in most force fields has been recognized as a potential problem since the early days of biomolecular simulations.^12,13^ Notable examples of situations where nonpolarizable models struggle involve the presence of high-charge-density ions influencing the structure of salt solutions,^14–16^ interactions of ions with lipids,^17,18^ interactions between charged amino acids,^19–21^ or the interactions between charged amino acids and acidic saccharides.^22^ Nonpolarizable force fields typically represent charge distributions by partial point charges located at the nuclei, yet this mapping of the electrostatic potential to partial charges is not unique. Moreover, partial charges remain constant during such simulations, thus explicitly excluding the description of electronic polarization effects. Another view of the problem is in terms of screening *via* the dielectric constant ɛ. Since classical MD keeps track of the motion of the nuclei, it naturally recovers the slow nuclear contribution to the dielectric constant (ɛ_nuc_) arising from molecular rearrangements. However, nonpolarizable MD fails by definition due to the use of fixed partial charges to capture the fast electronic polarization (ɛ_elec_), with this simplification allowing for atomistic simulations to reach microsecond or even millisecond time scales for large biomolecular systems.^23^

Importantly, electronic polarizability, while being non-negligible, is fairly constant in biological systems.^24^ For example, the values of ɛ_elec_ are 1.78 for pure water and 2.04 for hexadecane (mimicking the interior of membranes), with values for common salt and saccharide solutions, as well as more complex biological environments, falling within the same range.^24^ Electronic polarization is the dominant screening factor between charges in apolar or weakly polar environments. For example, the interior of a lipid membrane has ɛ ≈ 3, with contributions of ɛ_elec_ ≈ 2 and ɛ_nucl_ ≈ 1.5 (ɛ≈ ɛ_nuc_ ⇥ ɛ_elec_). In water, ɛ_elec_ ≈ 1.78 may seem at first sight small compared to the total polarizability of ɛ = 78. Nevertheless, the electronic component still leads to an additional attenuation of electrostatic forces to 1/1.78 ≈ 56% due to the roughly multiplicative effect of the nuclear and electronic polarizabilities.^25^ This missing electrostatic screening in nonpolarizable force fields thus often leads to overbinding and excessive aggregation of charged moieties in various biologically relevant environments.^26^ Electronic polarization can be accounted for explicitly in force field simulations *via* the introduction of atomic polarizabilities,^27^ fluctuating charges,^28^ or Drude oscillators,^29^ yet these approaches lead to a significant increase in computational cost. Moreover, as the polarizable models are typically not as excessively validated and fine-tuned as their nonpolarizable counterparts, they do not necessarily perform better in terms of accuracy.^30,31^ Two alternative strategies to account for the above-mentioned overbinding effects without explicitly introducing electronic polarizability are to modify the Lennard-Jones (LJ) potential or scale charges. CHARMM-based models typically opt for the former, applying additional repulsive terms in the LJ potential between selected atom types in order to prevent their association. While this heuristic approach, denoted as “NBFIX”,^32^ can fix specific overbinding issues, it also has severe shortcomings. Most importantly, as NBFIX is a modification of the Lennard-Jones potential, the response to external charges and electric fields cannot be properly captured in a physically well-justified way. Due to this fact, NBFIX may, for example, lead to repulsion between charged groups where association is actually observed experimentally. Another practical issue is that the repulsive NBFIX term needs to be derived separately for each involved pair of atom types.^24^ Finally, NBFIX could potentially hinder the avidity of molecular association relying on the specific coordination of charged groups.

The charge scaling approach suggested originally by Leontyev and Stuchebrukhov,^33,34^ accounts for electronic polarization in a mean-field way *via* the scaling of the partial charges. They called this approach “Molecular Dynamics in Electronic Continuum” (MDEC),^33^ with subsequent studies using the term “Electronic Continuum Correction” (ECC).^35^ As derived explicitly in the Methods section, charge scaling by a factor of ≈ 0.75 is mathematically equivalent to including the missing electronic part of the dielectric constant in the form of a dielectric continuum (*i.e.* 1.78 for water) into Coulomb’s law (Eq. (1)). Note also that the similarity of the ɛ_elec_ values for different biological environments justifies the use of a single fixed scaling factor.^24,34^ Within the ECC framework, a factor of ≈ 0.75 is thus used to scale all ionic charges.^24,33–35^

Within the last decade, our group has been extensively developing models based on the ECC approach, including force field parameters for monoatomic and molecular ions,^14,16,36–38^ proteins,^39,40^ and lipids.^41–43^ In parallel, other groups have applied charge scaling for simulations of ionic solutions,^44,45^ solid surfaces and their interfaces with aqueous solutions,^46–49^ biological systems^50^ and ionic liquids.^51–53^ Overall, there is growing interest in charge scaling models, which also signals the demand for a consistent and universal ECC-inspired force field,^35,54^ including parameters for biological macromolecules.^24,55^

Here, we present the first attempt for a consistent optimization patch of ECC-compatible models for biological systems based on the all-atom CHARMM36m/CHARMM36 force fields. Our model, abbreviated as `prosECCo75` (standing for “Polarization Reintroduced by Optimal Scaling of ECC Origin, scaling factor 0.75”, assumes the scaling factor of 0.75 for charges on ions and charged molecular groups. In this work, we demonstrate that `prosECCo75`, to a significant degree and in a physically justified way, cures overbinding artifacts related to interactions of aqueous ions, lipid membranes, amino acids, and monosaccharides. Employing `prosECCo75` improves the agreement in ion binding between simulations and experiments without compromising the description of biomolecular structures as following from the original CHARMM36m/CHARMM36 model. While charge scaling of simple ions has been addressed in our earlier studies,^14–16,36^ in this work, ECC is applied to zwitterionic and anionic lipids, essential amino acids, and acidic saccharides.

## 2 Methods

### 2.1 CHARMM36 Serves as a Starting Point for prosECCo75

We use CHARMM force fields as our templates since they are modular and provide a vast library of molecule types, which are updated in rolling releases. These include a protein model (“CHARMM36m”^56^) capable of reproducing the behavior of both structured and to some extent intrinsically disordered proteins,^57,58^ a vast library of lipids^59,60^ titled “CHARMM36”, and also parameters for mono- and polysaccharides introduced at a similar time also referred to as “CHARMM36”.^61,62^ Additional *ad hoc* repulsive interactions within the NBFIX concept have been regularly incorporated in the CHARMM force fields^32,60^ without the force field receiving a new version number. This leads to a somewhat unclear nomenclature. In this work, we differentiate between the CHARMM36/CHARMM36m model without any “NBFIX” parameters (here “`CHARMM36`”) and with all the current NBFIX additions (“`CHARMM36-NBFIX`”) that add specific nonbonded parameters to certain interactions between charged groups including ions, amino acids, proteins, lipids, and saccharides.

### 2.2 Introducing Electronic Polarization *via* Charge Scaling

Following the ECC approach,^33,34^ the missing electronic polarizability can be implemented in a mean-field way in MD simulations by scaling the (integer or partial) charges. This is evident when one writes the electrostatic interaction between two charged particles screened by the electronic polarization continuum as

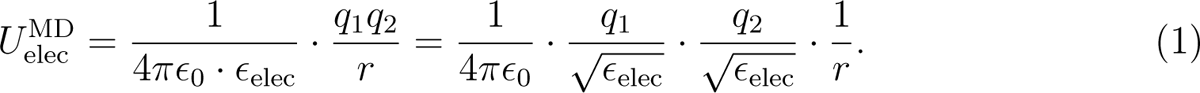

Here ɛ_0_ is the permittivity of vacuum, ɛ_elec_ is the high frequency dielectric constant arising from electronic polarization, q_1_ and q_2_ are the two atomic charges, and r is their interatomic distance. As seen in Eq. (1), electronic polarization screening is mathematically equivalent to scaling down the charges by a factor of ɛ*^−1/2^_elec_*, which equals to ≈ 0.75 in biologically relevant environments.

In our previous work, we introduced the ECC approach for Amber-based Lipid14 parameters for POPC (1-palmitoyl-2-oleoyl-sn-glycero-3-phosphocholine).^41^ While the scaling factors optimized for this Amber-based “ECC-lipids” model also improved `CHARMM36` simulations,^41^ Ca^2+^-binding affinity was still slightly overestimated. Moreover, we modified the Lennard-Jones σ parameters, which may have compromised the compatibility of lipid parameters when exposed to other molecules. Also, both changes (*i.e.* partial charges and Lennard-Jones) affect dihedral angles, which were originally optimized against experimental data in `CHARMM36`.^59^ Here, we avoid the above pitfalls by introducing ECC to `CHARMM36`. Our approach aims for minimal changes on interactions beyond the charge– charge ones without the need for *ad hoc* NBFIX corrections.

`CHARMM36` force fields are modular, meaning that molecules can be divided into smaller fragments, each with an integer charge, and these fragments serve as basic building blocks for all molecules. For example, in the zwitterionic POPC, the phosphate group has a total charge of –1, whereas the choline group has a total charge of +1. We strive to transfer this modularity to our ECC-corrected model to foster the transferability of the charge-scaled chemical groups. Therefore, we scale charges such that the absolute value of the total charge of any building block with an integer charge is reduced to 0.75 as mandated by ECC,^24^ corresponding to ɛ^water^_elec_ = 1.78. This scaling is applied only to blocks with non-zero charges, thus not affecting uncharged blocks. This minimal perturbation approach is important since the change of partial charges affects the dihedral angles, and we do not want to compromise the good description of structural ensembles by the `CHARMM36` model. Therefore, we do not modify the Lennard-Jones σ values in `prosECCo75`, contrary to our previous ECC-lipids work.^41^ Also, we apply changes to partial charges as far as possible from dihedrals critical, *e.g.*, for the conformations of protein backbone, lipid head groups, and saccharide ring puckering.

To demonstrate the validity of our approach, we present results for proteins and amino acids (scaling charges in termini and charged side chains); saccharides (scaling charges in carboxyl groups); and in membranes — phosphatidylcholines (PCs), phosphatidylethanolamines (PEs), and phosphatidylserines (PSs) — (scaling charges in their head groups). For proteins, the partial charges of carboxyl, ammonium, and guanidinium side chains and adjacent methyl groups bearing in total a charge of ±1 were uniformly scaled by a factor of 0.75. For the C-terminus, the scaled charges of carboxyl oxygens were taken from those of the side chains of the scaled aspartic and glutamic acids, with the charge of the carboxyl carbon adjusted to have the total charge of –0.75 on its block. For the N-terminus, the charges of the hydrogens in the NH^+^ group were taken from those of the lysine ammonium, and the charge of the nitrogen was adjusted for the group to have a total charge of +0.75. For acidic saccharides, the partial atomic charges of the carboxyl oxygens were taken from the acidic amino acids, while the charge of the carbon was adjusted to yield a total charge of –0.75. For lipids, we scaled the charges on phosphate oxygens so that the phosphate group has a charge of –0.75, while for choline the hydrogen charges were adjusted so that the total charge of the group is +0.75. The charges for secondary ammonium in PE and PS headgroups were taken from lysine amino acid side chain and further adjusted to have a total charge of 0.75. Section S1.1 in the Supporting Information (SI) contains a list of partial charges for all investigated lipids.

### 2.3 Simulation Parameters

All simulations were run using the default `CHARMM36`/`CHARMM36`m simulation parameters for GROMACS provided by CHARMM-GUI.^63^ We conducted all simulations in the isothermal-isobaric (NpT) ensemble, maintaining a temperature corresponding to experimental data with the Nośe–Hoover thermostat and a coupling time of 1 ps.^64,65^ The pressure was kept at 1 bar using the Parrinello–Rahman barostat with a 5 ps coupling time^66^ using a semi-isotropic scheme for membrane and osmotic pressure simulations and isotropic otherwise. The smooth particle mesh Ewald method was employed to calculate long-range contributions to electrostatics with a direct cutoff automatically adjusted around the input value of 1.2 nm.^67^ Lennard-Jones interactions were smoothly turned off between 1.0 and 1.2 nm using a force-based switching function.^68^ We kept track of atomic neighbours using buffered Verlet lists.^69^ We applied the SETTLE algorithm to constrain water geometry,^70^ with the P-LINCS algorithm constraining other covalent bonds involving hydrogen atoms.^71,72^

The lengths of almost all simulations reach at least 0.5 µs to provide sufficient statistics. The simulated systems employed either the original `CHARMM36`^56,59,62^ force field, or the newer variant incorporating NBFIX (`CHARMM36-NBFIX`),^60^ or the present `prosECCo75`. All simulations used the default `CHARMM36` TIP3P water (“mTIP” or “TIPS3P”)^73,74^ model unless stated otherwise. Additional parameters regarding the simulated systems are provided in sections S1.2 and S3.1 in the SI.

### 2.4 Osmotic Coefficients, Membrane-lipid C-H Bond Order Parameters, and Ion–Membrane Binding Isotherm and Residence Time

We calculated the osmotic coefficients, which are very sensitive to intermolecular interactions, from simulations using the method developed by Luo and Roux,^75^ which has been demonstrated to be an efficient tool for force fields refinement.^20,21,76,77^ The experimental reference data were either taken from the available literature (amino acids and polypeptides, collected by Miller et al. in Refs. 20 and 21) or measured by ourselves (monosaccharides, see below). The simulation values were obtained by restraining the solutes to a specific region of the simulation box using flat-bottom potentials and measuring the mean force hFi exerted by these solutes on the resulting semipermeable walls. Osmotic pressure was then calculated as П = 1/2 ⇥ hFi/A, where A is the cross-sectional area of the system. Finally, the molal osmotic coefficients were extracted as

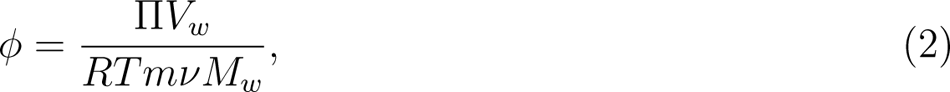

where V*_w_*is the partial molar volume of water (0.018 L·mol*^—^*^1^), R is the universal gas constant, T is the absolute temperature, m is the molality of the solution in the restrained part of the box, ⌫ is the Van’t Hoff coefficient (1 for neutral species and 2 for monovalent ions), and M*_w_* is the molar mass of water, 0.018 kg·mol*^—^*^1^.

We used C-H bond order parameters of lipids to evaluate membrane structure and ion binding affinity to membranes against experiments.^17,78^ These can be extracted from simulations as

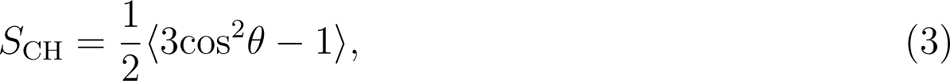

where ✓ is the angle of the C–H bond of interest with respect to the membrane normal. Importantly, the corresponding values can be measured with deuterium or ^13^C NMR, allowing a direct comparison between simulation and experiment.^78^

Because C-H bond order parameters are a good proxy for lipid conformational ensembles and membrane properties,^6,78,79^ we verified that the introduced charge modifications have only a minimal impact on membrane properties by calculating order parameters for all C-H bonds, including the headgroup, glycerol backbone and acyl chains, in membranes without additional ions. Results from simulations were compared with experimental datasets from the literature.^6,18,42,43,80^ Furthermore, changes in head group order parameters of α and β C–H bonds in phospholipids can be related to the number of bound cations in the membrane and used to evaluate the ion binding affinity in simulations against NMR experiments.^17,81^ To this end, we calculated order parameters from simulations with a defined added concentration of Ca^2+^ or Na^+^ ions for POPC and mixed POPC/POPS membranes. For these systems, we report the change in the order parameter, ΔS*^α/β^*, that is, the value of the order parameter at a given ion concentration minus the value for the system without additional ions. Ion concentration can be defined in two different ways: 1) System concentration refers to the concentration of all ions with respect to the total number of water molecules; 2) Bulk concentration is calculated from the number density of water molecules and the number density of ions in the bulk region (*i.e*., the region furthest away from the center of the membrane in the z direction). To report order parameters, we use the bulk concentrations as they correspond to the experimental concentrations. For cases (such as when evaluating density profiles) where we compare different force fields, we report system concentration as we compare runs for the same system with different potentials, resulting, in general, in different bulk concentrations.

Specifically for POPC lipids with Ca^2+^ ions, we also compare simulation results to the experimentally available binding isotherms, as obtained using atomic absorption spectroscopy.^82^ In simulations, an ion is defined to be bound to the membrane if its minimal distance from the nearest lipid oxygen atom is smaller than the 0.325 nm cutoff value set by the first minimum in the oxygen–cation radial distribution function (RFF).^83^ In addition, residence times of ions binding to membranes are reported as consecutive times for which the ion was closer than 0.325 nm to any lipid oxygen atom. The SI provides further details on all these simulation methods and analyses.

### 2.5 Experimental Measurements of Osmotic Coefficients

Osmolalities of saccharide–Na^+^ solutions were measured using a vapor pressure osmometer Osmomat 070 (Gonotec, Germany), following our established experimental protocol. ^84,85^ The osmometer was calibrated before each set of measurements with pure water and aqueous NaCl solutions. The osmolality of each solution was determined as an average of 10 readings. Details of the osmotic coefficient calculations from solution osmolality can be found in the SI.

### 2.6 Neutron Scattering Experiments on Ionic Solutions

We used neutron scattering techniques to measure the hydration shell around KCl, KBr, and KI ions in the solution and compare them with simulations. Heavy water (99.9 atom % D) and light water (H, 18 Mω) were mixed together (78.688 g H_2_O and 48.975 g D_2_O).

The hydrogen in this mixed water had an average coherent neutron scattering length of 0 fm (*i.e.*, for this mixture, the scattering from hydrogen and deuterium cancel each other). KCl, KBr, and KI were dried in a vacuum oven at 150*^○^*C overnight. 4 M solutions of potassium halides were then prepared by direct dissolution of salt in water. In each case, 5 mL samples were prepared. From each solution and null water, 0.75 mL was transferred to a null scattering Ti/Zr cell, and neutron scattering data of each sample were recorded on the D_2_O diffractometer for around two hours. The scattering data was then corrected for multiple scattering and absorption prior to being normalized versus a standard vanadium sample to yield the total scattering pattern for each solution.

To characterize these solutions, we use a technique similar to that used in our previous work.^86^ Null scattering water solutions have a large incoherent background that mostly scales with the atomic concentration of ^1^H in the solution. Subtracting the total scattering patterns of two null scattering solutions mostly cancels out this background and makes subsequent analysis simpler. If the total scattering pattern of null water is directly subtracted from that of a potassium halide solution, the residual also largely cancels out the oxygen–oxygen correlation (S_OO_), which constitutes around two-thirds of the total coherent scattering from these solutions. The leftover contribution contains valuable information regarding ions and their surroundings and ion pairing, and the details on the different contributions for each system can be found in the section S4.3 in the SI. This leftover signal can also be directly compared to results from simulations.

## 3 Results

In the following subsections, we demonstrate how the inclusion of electronic polarization by charge scaling significantly improves the interactions between ions and various classes of charged biomolecules without compromising the biomolecular structures reproduced well already by the original `CHARMM36` force field.

### 3.1 prosECCo75 Provides Realistic Binding of Ions to Lipid Membranes

Here, we rely on a direct comparison with the experiment to validate the charge scaling approach in lipids and ions. We verify the ECC approach on membranes composed of phosphatidylcholine — POPC and 1,2-dipalmitoyl-sn-glycero-3-phosphocholine (DPPC), which are two common and extensively studied zwitterionic lipids. As PS is the most abundant charged lipid type in the mammalian plasma membrane,^87^ we also include 1-palmitoyl-2-oleoyl-sn-glycero-3-phospho-L-serine (POPS) in our study. While we mostly focus on the headgroup response to surrounding ions (vide infra), we also evaluate how well the lipid model reproduces structural properties in the absence of ions (see SI). Also, for 1-palmitoyl-2-oleoyl-sn-glycero-3-phosphoethanolamine (POPE) and cholesterol, we benchmark their behavior in the absence of ions. Below, we present results for all studied membranes using three different force field models: 1) the original version of `CHARMM36`, 2) its variant `CHARMM36-NBFIX`, and 3) our `prosECCo75`, along with experimental data wherever available.

As shown in Figure 1, the `prosECCo75` implementation for PCs significantly reduces the binding of Ca^2+^ ions to POPC membranes when compared with `CHARMM36`, yielding results in line with experiment.^82^ `prosECCo75` provides an overall significantly better agreement with the experiment, namely, it captures the effect of an increasing ion concentration while slightly undershooting the number of bound ions. For `CHARMM36`, a significant Ca^2+^ density is found around the phosphate group, while almost no cations remain in bulk water (Figure 2b). For `prosECCo75`, the binding is significantly reduced. An even smaller number of bound ions is observed for `CHARMM36-NBFIX`, yet the difference from `prosECCo75` in density profiles is small (CaCl_2_ panel in Figure 2b). Interestingly, this small difference in density profiles corresponds to a major change in the number of bound ions (Figure 1). This effect results from the character of the NBFIX potential,^32,88^ which strongly repels Ca^2+^ ions from the membrane interface.

**Figure 1:**
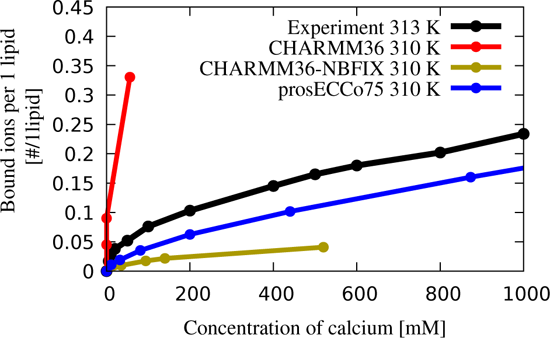
Binding isotherm of Ca^2+^ ions to POPC membrane. The concentration of Ca^2+^ ions is reported as the bulk concentration. Bound ions are defined by a 0.325 nm cutoff from either phosphate or carbonyl oxygen atoms, corresponding to the first minimum in the radial distribution function. The result is rather insensitive to the exact value of the cutoff.^83^ The error estimates for the values calculated from simulations are smaller than the size of the markers. The corresponding atomic absorption spectroscopy data were taken from Ref. 82.

**Figure 2:**
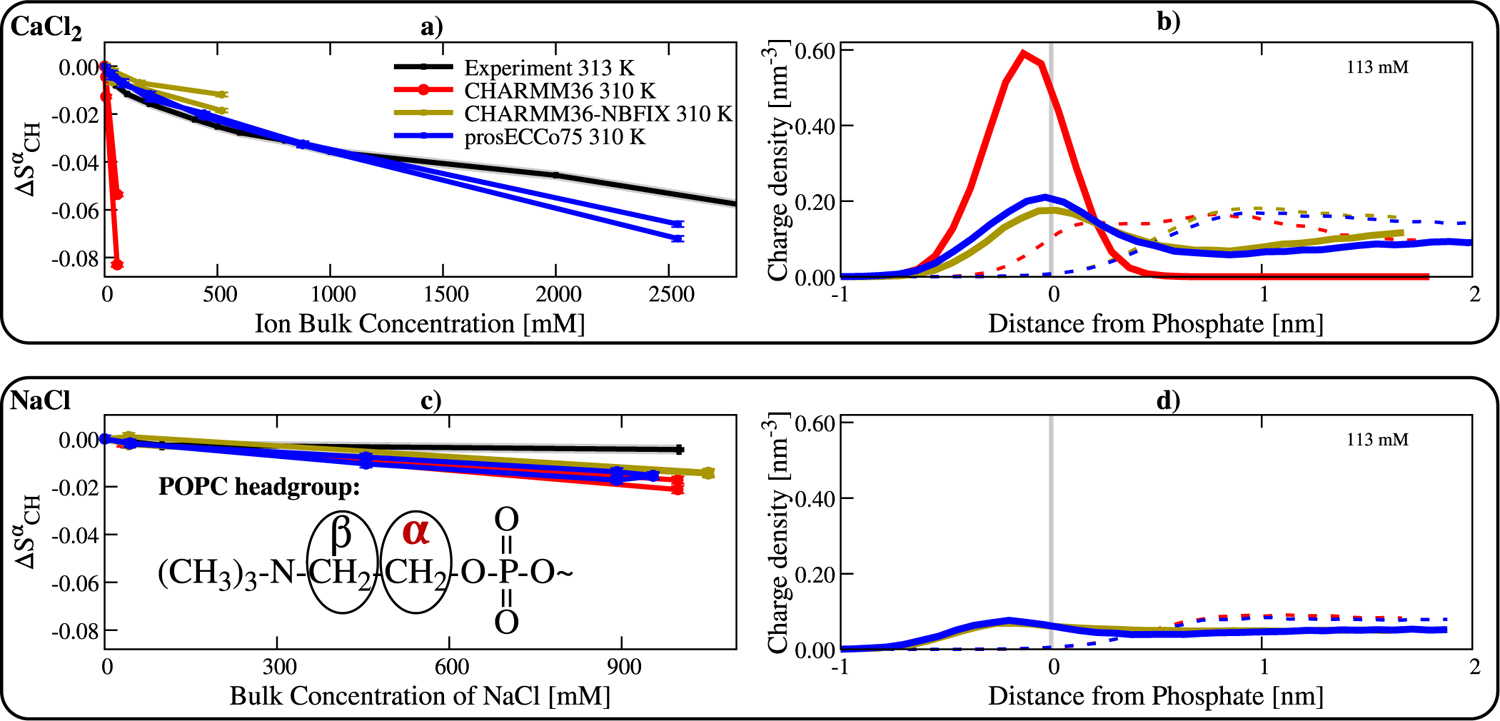
Binding of Ca^2+^ and Na^+^ ions to POPC membranes. Top panel: Ca^2+^ ions. Bottom panel: Na^+^ ions. Panels a) and c) show the order parameter response on the α position of POPC lipids as a function of bulk salt concentration. Experiments are from Ref. 82. Panels b) and d): density profiles of Ca^2+^ and Na^+^ ions calculated from MD simulations (shown by solid lines), as well as the Cl*^—^* counterions (dashed lines), are shown for the three different models. All the density profiles are calculated along the membrane normal and centered around the maximum density of the lipid phosphorus atoms (vertical grey line). The simulation error bars are smaller than the size of the markers.

We further benchmark `prosECCo75` for phospholipids using the electrometer concept.^17,81^ Namely, we compare the responses of the lipid headgroup order parameters to increasing salt concentration in simulations with those measured by solid-state NMR. When the headgroup order parameters fit the experimental values well, the responses of headgroup order parameters to ions relate directly to the adsorption of these ions to the headgroup region. We see that `CHARMM36` significantly overestimates the response of the order parameter in the α position (ΔS*^α^*) in a POPC membrane to increasing Ca^2+^ concentration (Figure 2a). This well-known deviation results from a significant Ca^2+^ overbinding^17,18^ as seen in Figure 2b. Incorporating NBFIX^32,88^ overcorrects this effect, thus leading to an overly too weak response. `CHARMM36-NBFIX` lacks accumulation and even displays depletion of Ca^2+^ at the interface, especially for larger ion concentrations, see Figure S13 in the SI. The NMR order parameter data (ΔS*^α^*),^82^ in agreement with the binding isotherm,^82^ support the observation that the strength of Ca^2+^ ion binding to POPC membranes is significantly improved in the `prosECCo75` force field as compared to `CHARMM36-NBFIX` (Figure 2a). Only at very high Ca^2+^ concentrations (> 1 M CaCl_2_), `prosECCo75` slightly deviates from experiment toward overbinding (Figure S12). Interestingly, Ca^2+^ density profiles at concentrations above 500 mM show lower accumulation of Ca^2+^ at the interface compared to the bulk (Figure S11a). The clear improvement for `prosECCo75` over `CHARMM36-NBFIX` shown by the binding isotherms and ΔS*^α^* exists despite the differences in the Ca^2+^ and Cl*^—^* density profiles being small in general, see Figure 2b. Additionally, a comparison of head group responses of DPPC and POPC to CaCl_2_ is provided in Section S1.5.7 in the SI.

For all the tested force fields, the experimental response of head group order parameters (ΔS*^α^*) to increasing Na^+^ concentration is reasonably well reproduced yet slightly overestimated (Figure 2c). The responses of `CHARMM36-NBFIX` and `prosECCo75` fit the experiment only marginally better than that of `CHARMM36`, which is primarily due to the fact that there is only a small Na^+^ accumulation in the headgroup region (Figure 2d). Also, Na^+^ density profiles for the three models show only minor differences. Densities for Na^+^ at the membrane at varying concentrations can be found in Figure S11b in the SI. Overall, our results indicate that Na^+^ may slightly overbind to POPC membranes in `prosECCo75`, nevertheless, with a consistently low Na^+^ binding to the membrane for the whole concentration range (Figure S12).

The comparison between Ca^2+^ and Na^+^ responses is not straightforward. While Ca^2+^ binding is systematically larger than that of Na^+^ for POPC membranes at biologically relevant concentrations (Figures S12 and S13), the differences in surface densities between these two ions are small, particularly at large concentration. However, when considering charge densities (Figures 2b and 2e), the difference between the two cations is much more evident, particularly at low, more biologically relevant Ca^2+^ concentrations, where Ca^2+^ binds significantly more to POPC than Na^+^.

The scaling of the lipid charges in the head group region affects not only the strength of ion binding but also the molecular details of the ion binding modes (Figure S10 in the SI) and the residence times (Figure 3) of Ca^2+^ in membranes. For `CHARMM36`, all the Ca^2+^ ions that bind to a membrane stay bound for the remaining duration of the simulation. Thus, for most of these ions, we observe residence times longer than 1 µs (left panel in Figure 3). The situation is very different in the case of `CHARMM36-NBFIX` and `prosECCo75`, where we observe in both cases numerous binding and unbinding events throughout the simulations. With `CHARMM36-NBFIX` and `prosECCo75`, the residence times are up to 5 ns and 63 ns, respectively. Experimentally, an upper bound for this fast Ca^2+^ exchange process is set by IR to 150 ns^89^ or by NMR spectroscopy to 10 µs.^82^ This upper bounds are consistent with both `CHARMM36-NBFIX` and `prosECCo75`, but not with `CHARMM36`.

**Figure 3:**
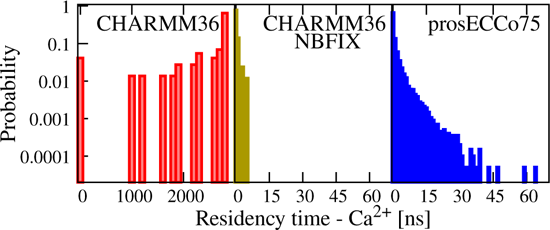
Ca^2+^ residence times in the POPC membrane. Residence times are calculated as a consecutive time for which the given ion is within the cutoff of 0.325 nm from any lipid oxygen. All the values are calculated from simulations with 450 mM of CaCl_2_. Unbinding events are absent in `CHARMM36` simulations, *i.e.*, the reported binding times correspond to the difference of the total simulation time (3 µs) and the time at which a Ca^2+^ binds.

Ca^2+^ ions also prefer to form complexes with a larger number of lipids when using force fields where the ion binding is stronger (Figure S10 in the SI). Namely, in `prosECCo75`, Ca^2+^ mainly binds to one lipid (≈ 50%), but often it complexes two (≈ 35%) or even three lipids (≈ 10%) while in `CHARMM36-NBFIX` it mostly binds to one lipid (≈ 75 %). These coordination numbers agree reasonably well with those resulting from fitting simple binding models to experiments.^82^ In contrast, results for `CHARMM36` are different, resulting in larger complexes, *i.e.* 3–4 lipids per Ca^2+^, which form large aggregates as reported previously.^83^ We also observe that the preferred binding sites of Ca^2+^ (*i.e.* phosphate versus carbonyl oxygens) vary between models (Table S12 in the SI). Unfortunately, there are no experimental data to directly compare to.

Another lipid for which the response of the head group order parameters to Ca^2+^ concentration was experimentally measured is POPS in POPC:POPS mixtures at a ratio of 5:1. This mixture is used as the addition of Ca^2+^ to pure POPS leads to the formation of precipitates.^90,91^ The POPC and POPS headgroup responses in the 5:1 mixture to CaCl_2_ are shown in Figure 4. A substantial improvement with respect to experimental data, in particular for the β carbon, is observed with `prosECCo75` compared to `CHARMM36` and `CHARMM36-NBFIX`.

**Figure 4:**
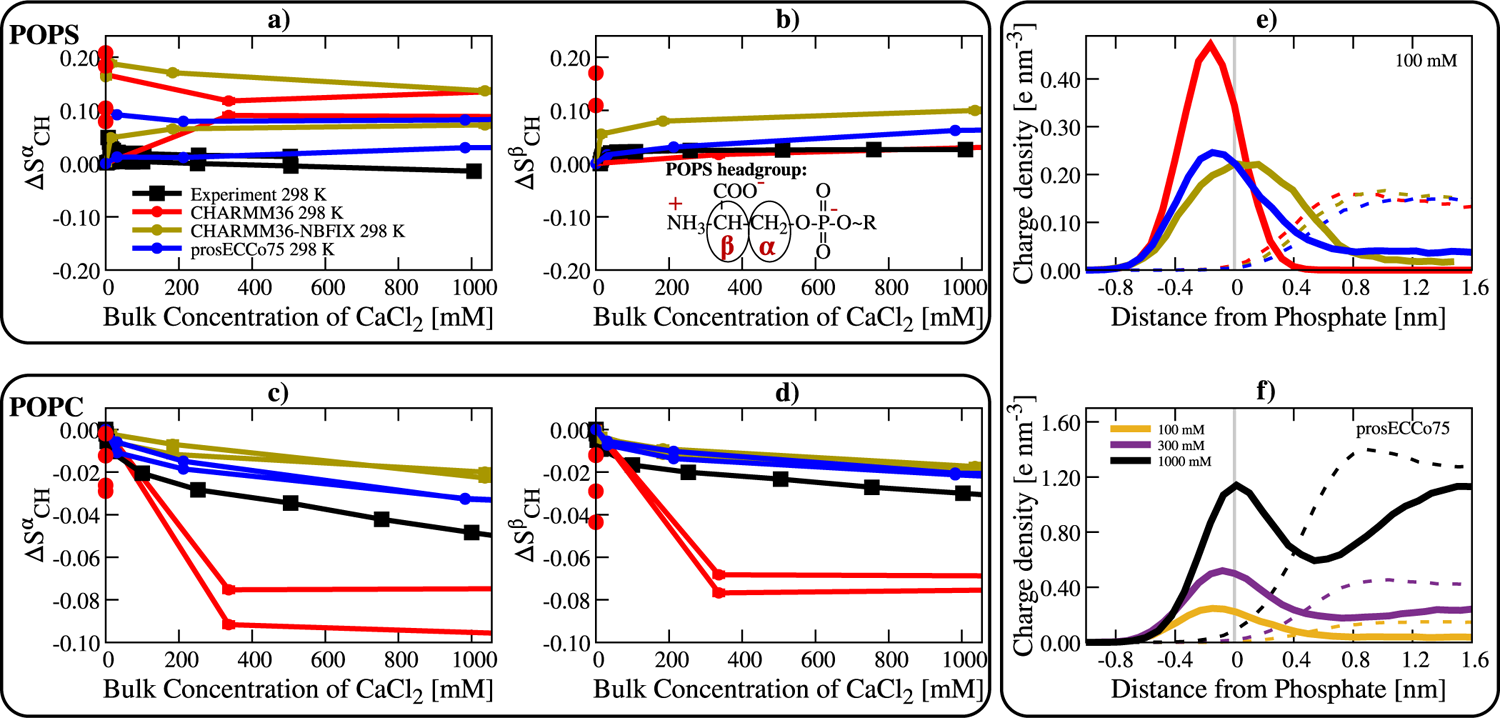
Binding of Ca^2+^ ions to 5:1 POPC:POPS mixture membranes. Panels a) and b): behavior of POPS in the mixture. Panels c) and d): behavior of POPC in the mixture. All these panels show the response in a lipid bilayer as a function of the bulk concentration of Ca^2+^ in the system. Panel e) compares the calcium density profiles centered around maximum phosphate density for the three force fields and panel f) shows the calcium density profiles of `prosECCo75` model at three different concentrations. Na^+^ was used to neutralize POPS charges. Experimental data are from Ref. 91. In the case of `CHARMM36`, the multiple points at a bulk concentration of 0 mM result from all Ca^2+^ ions being bound to the lipids in several systems with different total numbers of Ca^2+^ per system.

In `prosECCo75`, results for one of the hydrogen atoms attached to the α carbon match the experimental line. However, the splitting between the two hydrogens is overestimated compared to the experiment. Both `CHARMM36` and `CHARMM36-NBFIX` exaggerate the ion effect for both hydrogen responses. This is anticipated for `CHARMM36` POPS as it shows a significantly larger binding to Ca^2+^ than `prosECCo75`. But even for `CHARMM36-NBFIX`, which exhibits a similar binding affinity of Ca^2+^ for POPS as in `prosECCo75`, the head group order parameter response is significantly different from the experiment. It is important to mention that the absolute order parameters of the POPS head group in the absence of additional ions are somewhat off from the experimental values for all investigated force fields (Figure S7 in SI). Moreover, the simulated order parameter response (ΔS*^α^*) to ions does not necessarily even qualitatively follow the experimental trend (Figures S14, S15 and S16). Overall, `prosECCo75` provides a somewhat better agreement with the experiment than `CHARMM36`, yet our data indicate that there is a need to refine not only the PS–cation interactions (*e.g.*, Na^+^ clearly overbinds) but also the PS head group itself. The response of POPC in the 5:1 POPC:POPS mixture is also slightly improved with `prosECCo75` over `CHARMM36-NBFIX`, which is already a drastic improvement from `CHARMM36` (Figures 4c and 4d). Based on the above comparison with the NMR data we can conclude that `prosECCo75` captures ion binding to membranes significantly better than `CHARMM36` and, even more importantly, it also shows improvement over `CHARMM36-NBFIX`, likely due to its a physically better-justified foundation.

While the response to ions is improved in `prosECCo75` over the `CHARMM36` and `CHARMM36-NBFIX` models, at the same time, the already good description of the membrane structure should not be compromised by the changes made in `prosECCo75`. To compare the structures for POPC, DPPC, POPS, POPE, and cholesterol-containing membranes produced by `prosECCo75` and `CHARMM36`/`CHARMM36-NBFIX` models, we compute order parameters of the head groups and acyl chains, form factors, transition temperatures, and areas per lipid. These values are compared to experiments in section S1.5 in the SI. Our results show that in the absence of ions, `prosECCo75` agrees equally well with the experiment in essentially all calculated properties as `CHARMM36`/`CHARMM36-NBFIX`. (Note that in the absence of ions, `CHARMM36` and `CHARMM36-NBFIX` are identical for lipids, except for certain POPS counterions interactions.)

### 3.2 prosECCo75 Reduces Excessive Protein–Protein Interactions

Osmotic coefficients provide information about intermolecular interactions among solute molecules. Smaller values indicate that the solutes tend to aggregate, imposing a reduced osmotic pressure on a semipermeable membrane. The osmotic coefficients extracted from simulations using common protein force fields are generally too low compared to experiments, indicating an excessive attraction among amino acids.^20,21^ Earlier attempts to correct this discrepancy were based on empirical scaling of the Lennard-Jones parameters (similarly to NBFIX), which led to a significant improvement.^20,21^ However, there is no good physical justification for this tuning. Since single amino acids are zwitterionic and some even charged, it seems reasonable to assume that electrostatic interactions largely dominate their interactions at high concentrations used in osmotic coefficient measurements, thus `prosECCo75` may provide an improvement over `CHARMM36-NBFIX`. To verify this assumption, we calculated the osmotic coefficients for all essential amino acids, as well as for some short polypeptides, at varying concentrations. These osmotic coefficients of all the studied solutions are compared to the experimental values in Figure 5.

**Figure 5:**
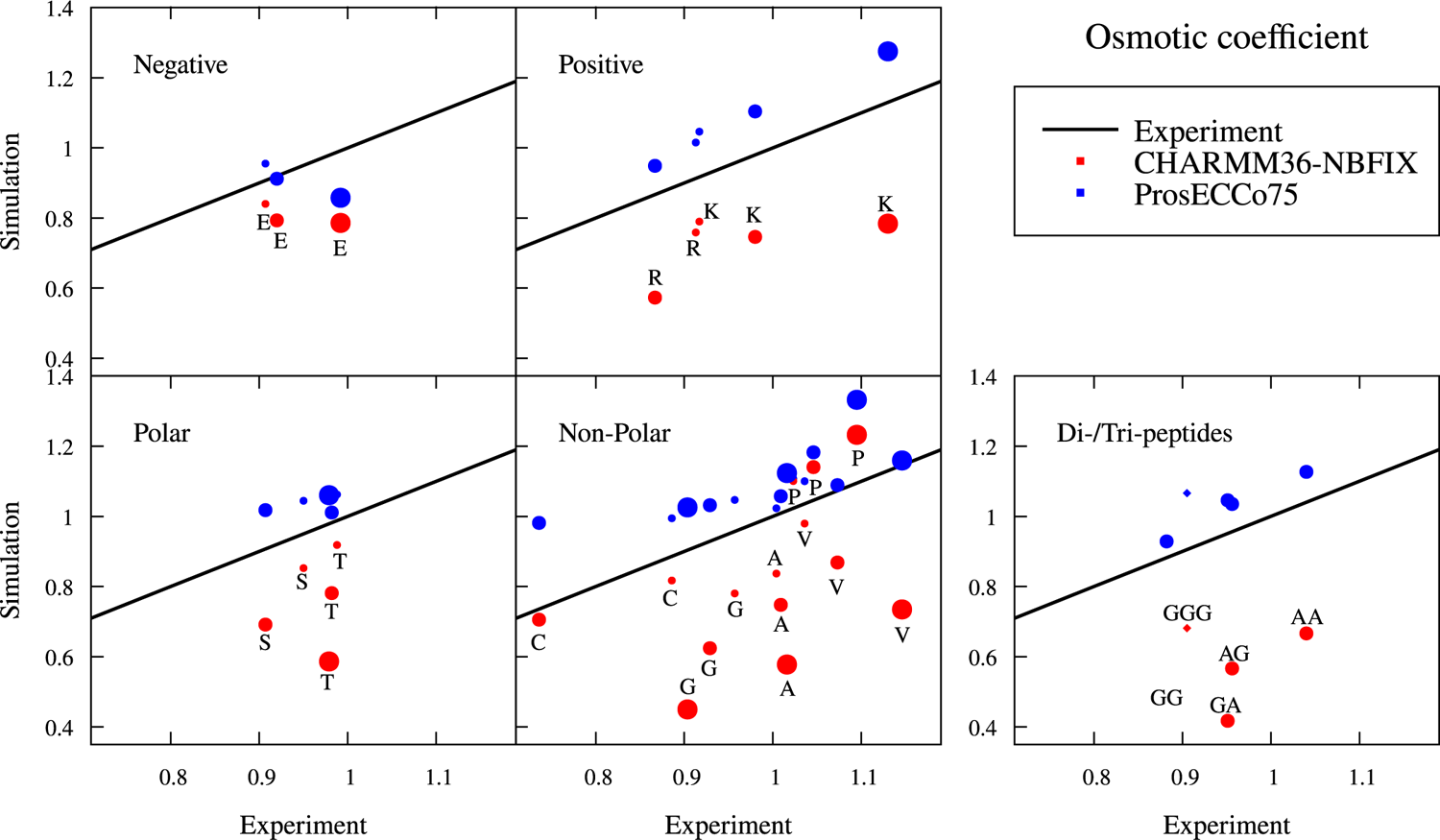
Interaction between amino acids, dipeptides, or tripeptides in solution. Comparison of osmotic coefficients of amino acid, dipeptide, and tripeptide solutions between experiment and simulations. Na^+^ counter ion is used for charged amino acids. Larger values mean less attraction between amino acids, dipeptides, and tripeptides. The increasing size of the symbol indicates 0.5 M (except tripeptide 0.3 M), 1 M, and 2 M solutions, respectively. Tripeptide points are shown with the diamond symbol. Simulation errors are 0.02, given as the standard deviation.

We see that `prosECCo75` provides a significant improvement in osmotic coefficients over `CHARMM36-NBFIX` for all amino acids and small polypeptides. Still, the `prosECCo75` values show a small but systematic overestimation, indicating that the simulated amino acids are slightly less “aggregated” than they should be. The amino acids for which experimental data exist are classified into four categories (anionic, cationic, polar, and apolar), and `prosECCo75` provides a better agreement with an experiment than `CHARMM36-NBFIX` for all these categories. There seems to be no systematic correlation between amino acid concentration and the quality of agreement with the experiment, suggesting that the `prosECCo75` approach is rather universal. Similarly, the behavior of dipeptides (AlaAla, GlyGly, AlaGly, and GlyAla) and a tripeptide (GlyGlyGly) with apolar side chains suggests that the scaling of atomic charges on both termini has a major impact on the osmotic coefficients. Finally, a critical aspect of protein simulations concerns the backbone dihedrals, which ultimately define the secondary and tertiary structures of a protein. Importantly, these dihedrals might be affected by the tempering with the charges. However, our charge scaling involves only the far side-chain charged groups, largely avoiding such a problem (Figure S20). Furthermore, simulations of some structurally challenging intrinsic disorders proteins show only differences within the calculated error ranges for `prosECCo75` and `CHARMM36-NBFIX` (Figures S21 and S22). Overall, `prosECCo75` provides a clear improvement in the description of interactions between amino acids — including the uncharged ones — by decreasing the excessive electrostatic interactions of `CHARMM36-NBFIX`.

### 3.3 prosECCo75 Tones Down Excessive Saccharide–Saccharide Interactions

Moving on to saccharide-containing species, we considered here two acidic saccharides, D-galacturonic acid and D-glucuronic acid, which are the oxidation products of galactose and glucose and are common compounds in glycosaminoglycans, pectin, and gums. Moreover, these acidic saccharides are charged, unlike their non-oxidized counterparts, providing an excellent test case for `prosECCo75`. The osmotic coefficients of these acidic saccharides as a function of their concentration from simulations with different force fields are compared to the results of our experiments in Figure 6.

**Figure 6:**
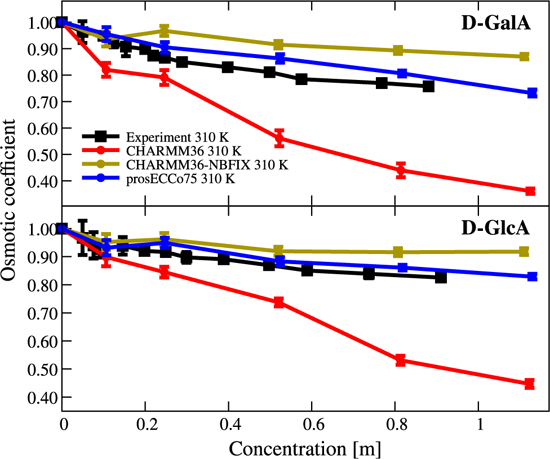
Interaction between charged monosaccharides in solution. Comparison of osmotic coefficients of charged monosaccharide solutions between experiment and simulations. Larger values mean less attraction between solutes.

From Figure 6, it is evident that the observed trend for the acidic saccharides is similar to the amino acids in Figure 5. With increasing concentration, `CHARMM36` with unscaled charges overestimates the intermolecular attraction, which leads to significantly lower osmotic coefficients as compared to the experiment. Once the charges of the saccharide carboxyl groups and the Na^+^ counterions are scaled in `prosECCo75`, the electrostatic attraction is no longer excessive. NBFIX has a similar but too strong effect, *i.e.*, it inhibits ionic pairing more than required to match the experiments leading to noticeably overestimated osmotic coefficients. Unlike the other force fields, `prosECCo75` thus provides an excellent agreement with the experiment over the investigated concentrations, only slightly over-correcting the association tendency, as indicated by a bit higher-than-experiment osmotic coefficients, particularly at intermediate concentrations of the D-galacturonic acid. Our results support the application of `prosECCo75` as an improvement to `CHARMM36` in modeling charged saccharides, especially when modeling the biologically important glycosaminoglycans such as heparan sulfate and hyaluronic acid, with alternating uronic acids and amino saccharide.

### 3.4 ECC Ions are Required for Biomolecular Simulations Using prosECCo75

Biological aqueous environments are enriched in salt ions. These ions play critical roles in signaling pathways, in balancing the osmolarity between biological environments, and in the creation of potentials across the membranes required for cell homeostasis. Scaled-charge force fields for biomolecules require compatible ions, *i.e.* they must have similarly scaled charges (by 0.75 in the case of `prosECCo75`).^24^ Here, we append Br*^—^* and I*^—^* anions to our list of available ions with scaled charges: Na^+^ (Na s^14^), Li^+^ (Li s^16^), K^+^ (K s^92^), Ca^2+^ (Ca s,^36^ Ca 2s^15^), Mg^2+^ (Mg s^93^), and Cl*^—^* (Cl s,^36^ Cl 2s^16^). In the previous sections of this work, we used the Na s model^14^ for Na^+^, the K s model^92^ for K^+^, the Ca 2s model^15^ for Ca^2+^ except POPC/POPS mixtures where Ca s performed significantly better,^36^ and the Cl 2s model^16^ for Cl*^—^*.

We parameterized the missing Br*^—^* and I*^—^* anions (Figure S24) using densities and structural data from neutron scattering. For Br*^—^*, we produce two models, Br s and Br 2s. This follows Cl*^—^*, which possesses two variants Cl s^36^ (better density) and Cl 2s^16^ (better agreement with structure, *i.e.*, neutron scattering experiments). With I*^—^*, one model could match the density and structural experimental data for the “ s” and “ 2s” series while preserving their differences.

These two new halide ions (Br*^—^* and I*^—^*) provide reasonable densities^94^ in combination with K s at physiologically relevant conditions when using the SPC/E water model^95^ (pure `CHARMM36` TIP3P water^73,74^ does not have a reasonable density), see Table S22.

We can further benchmark our new ion models using the radial distribution function between “all” atom pairs, which is available *via* neutron scattering for KCl, KBr, and KI. As the water signal is overwhelming, to best characterize RDF (G) involving ions in neutron scattering experiments, the hydrogen contribution can be removed using null water^86^ and the oxygen–oxygen contribution by subtracting that of pure water signal. The leftover signal mainly contains the cation oxygen solvation shell, the halide anions oxygen solvation shell, and the ion pair contributions, which better characterize the salt solvation shell. We experimentally measured such RDFs, which can also be directly computed from our simulations for comparison (Figure 7 and Figure S25 for the “ 2s” and “ s” K^+^A*^—^*anion series, respectively).

**Figure 7:**
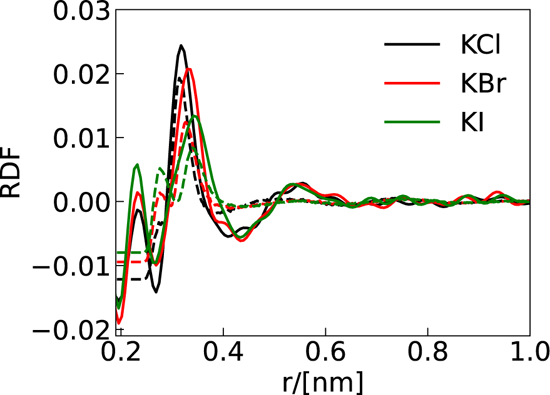
Solvation shell of potassium chloride (KCl), potassium bromide (KBr), and potassium iodide (KI). The solid lines are the experimental neutron scattering results and the dashed lines are the simulation results using K s, Cl 2s, Br 2s, and I s parameters.

The source of the negative RDF values signals that the oxygen–oxygen scattering patterns for pure and salt solutions are not equal due to the ions distorting the hydrogen bond network and hence, changing the oxygen–oxygen distribution.

When using the “ 2s” and “ s” K^+^X*^—^* anion series, we obtain reasonable agreement for the g_X_*—*O and g_K_+X*—* (r) pointing to adequate sizes of the halides respect to their environment and the ion pairs. Therefore, one should expect more reasonable differences and selectivities when comparing halides in biological systems.^50^

## 4 Discussion and Conclusions

The present results demonstrate that `prosECCo75` provides an improved agreement with experiment when charge–charge interactions between biomolecules (lipid membranes, amino acids/proteins, and saccharides) and ions are central to the measured property. At the same time, we have shown that `prosECCo75` does not deviate significantly from `CHARMM36-NBFIX` in cases where charge–charge interactions play only a minor role. In addition, the previous success of the ECC approach in a broad range of applications and with other underlying force fields suggests that charge-scaling is transferable at least to some degree.^37,39–42^

Still, it would be unrealistic to expect that inaccuracies of the underlying force fields can all be fixed via charge scaling. For example, the measured POPC or DPPC headgroup order parameter for `prosECCo75` as a function of Ca^2+^ concentration always follows the experimental line measured for POPC while the Amber-based ECC models follow the DPPC line instead, see Figure S18. It is unclear why experiments and simulations provide different results in this case. For the charged PS headgroup, the situation is even more complicated because the binding details are highly sensitive to the underlying force field parameters.^18^ Therefore, ECC-based force fields have problems correctly capturing the response of order parameters to cation binding as shown in Figure 4 and, for different force fields, in Refs. 42,43. Nevertheless, the response of PS headgroups is improved in `prosECCo75`, suggesting that also binding details of Ca^2+^ to PS is more realistic, although we still observe a modest sodium overbinding, see Figure S1.5.8 in the SI. Moreover, some of the underlying force field issues can be further emphasized by charge scaling. For example, when comparing ECC-Amber-based and `prosECCo75` force fields for polyanionic peptides, these peptides condense differently in the presence of counterions.^39,96^ Also, different implementations of the ECC protocol can lead to quantifiable differences (See Section S1.5.6 in the SI), justifying the charge optimization performed in this work. Overall, the observed differences between various ECC implementations are rooted in subtle differences in the underlying force fields. In particular, most nonpolarizable models of neutral polyatomic molecules implicitly deal with the effects of electronic polarization to some extent, as their partial charges are often fine-tuned to reproduce an experiment, which may lead to “overscaling” when applying ECC. Further improvements thus require a more rigorous *de novo* force field development starting with a water model fully compatible with the ECC concept.^97^

It is a legitimate question whether `prosECCo75` presents a substantial improvement over `CHARMM36-NBFIX` and hence justifies another revision of the original force field. Indeed, both approaches lead to a similarly reasonable agreement with experiments in most cases. However, in the case of `CHARMM36-NBFIX`, this agreement results from the *ad hoc* repulsive potential between the cations and the lipid headgroups. While this can often improve the agreement with the experiment, it can also result in undesired side effects in some cases. As an example, the repulsion preventing the overbinding of cations to POPC bilayers also leads to their unnaturally low levels at the membrane surface (Figure 1). This, in turn, leads to an inadequate description of biological processes where ion-coordinated binding is important, such as the membrane anchoring of the C2 domain of protein kinase C α unit (PKCα). The binding of the anionic loops of PKCα-C2 to a phosphatidylserine lipid is bridged by Ca^2+^,^98^ and thus capturing this binding mode requires properly balanced ion–protein and ion–lipid interactions. As we demonstrated recently, PKCα-C2 binding mode involves the bridging by Ca^2+^ with `prosECCo75`, in line with the crystal structure (PDB:1DSY^98^).^24^ In contrast, `CHARMM36-NBFIX` results in the adsorption of the protein at an incorrect orientation. In the absence of NBFIX, the PKCα C2 domain does not even adsorb to the PC/PS membrane, as the excessive ion–protein and ion–lipid binding render both the protein and the membrane effectively highly cationic and thus mutually repulsive. In addition to the inability to model such detailed binding modes, the NBFIX approach also requires a significant amount of parameterization work, as the repulsive terms must be adjusted separately for the different pairs of atom types. Also, the ECC correction does not always result in repulsion. For example, applying the ECC approach to small charged organic molecules can increase their binding strength to membranes in qualitative agreement with experiment,^37^ where an NBFIX approach might not be able to accommodate such increase of binding or the very least be too specific. This workload of tuning the NBFIX parameters can be significantly reduced by applying as an alternative the physically sound and universal `prosECCo75` approach.

Although the scaling factor for charges within the ECC framework is dictated by the medium’s electronic polarizability and should, thus equal 0.75 in aqueous solutions, other scaling factors have also been used in the literature. For example, early lipid bilayer simulations recognized that charge scaling could compensate for the lack of electronic polarization. Yet, these studies used a somewhat arbitrary scaling factor of 0.5.^12,99,100^ In contrast, for ionic solutions, a larger scaling factor of 0.85 was shown to provide an excellent agreement with experiments.^44,45^ Furthermore, benchmark ab initio MD simulations showed recently that while the scaling factor prescribed by ECC correctly describes the long-range interaction, a somewhat weaker scaling factor of ≈ 0.8 better captures the short-range direct interaction between charged ions.^101^ These results may explain why — despite the significant improvements presented here — a small but systematic under-binding is observed for the `prosECCo75` force field in many of the applications presented here. This may indicate that in future development, the charge scaling factor may be slightly adjusted upwards.

To conclude, we showed that the inclusion of electronic polarization in a mean-field way via charge scaling into the `CHARMM36` force field results in an improved description of the electrostatic interactions in applications involving interactions of ions with biomolecules. In particular, we highlight here improvements in ion binding to lipid bilayers and in the intermolecular interactions between charged saccharides and amino acids. The present model, denoted as `prosECCo75`, is based on modifying partial charges of atoms in a way that maintains the building block nature of `CHARMM36` with the molecular fragments whose charge was originally an integer number being scaled by a factor of ±0.75. In addition, the universality of the ECC framework streamlines the adaptation of `prosECCo75` to the ever more complex biomolecular ensembles for MD simulations. It is our aim that the significant improvements presented here, along with the portability of the fragment approach, inspire other researchers to adopt `prosECCo75` in their work.

## Acknowledgement

M.R.-F. and H.M.-S. acknowledge the support of the Czech Science Foundation (project 19-19561S). M.R.-F. and V. C. C. acknowledge the support from the Charles University in Prague and the International Max Planck Research School in Dresden. M.J. acknowledges support from the Research Council of Finland (grant no. 338160) and the Emil Aaltonen foundation. P.J. acknowledges support from the European Research Council *via* an ERC Advanced Grant no. 101095957. R.N. and O.H.S.O acknowledge the support from the Research Council of Finland for funding (grant no. 315596, 319902, and 345631) and CSC – IT Center for Science for computational resources. R.N acknowldeges financial support from the Emil Aaltonen Foundation.

## Supporting Information Available

Detailed simulation methods. Results demonstrating that `prosECCo75` does not compromise the structural ensembles of the `CHARMM36` force field. Additional results supporting the findings of the main text.

## Notes

### Competing Interest Statement

The authors have declared no competing interest.

