## Supplementary information for "Effective Inclusion of Electronic Polarization Improves the Description of Electrostatic Interactions: The prosECCo75 Biomolecular Force Field"

### Contents

**S1 Supplementary Information Membranes** **S-4**

S1.1 Partial Charges for Lipids . . . . . . . . . . . . . . . . . . . . . . . . . . . S-4

S1.2 Simulated Lipid Bilayer Systems . . . . . . . . . . . . . . . . . . . . . . . . . S-7

S1.2.1 POPC . . . . . . . . . . . . . . . . . . . . . . . . . . . . . . . . S-7

S1.2.2 DPPC . . . . . . . . . . . . . . . . . . . . . . . . . . . . . . . . S-9

S1.2.3 POPS and POPS/POPC Systems . . . . . . . . . . . . . . . . . . . S-9

S1.2.4 POPE Systems . . . . . . . . . . . . . . . . . . . . . . . . . . . . S-11

S1.2.5 Systems with Cholesterol . . . . . . . . . . . . . . . . . . . . . . . S-12

S1.3 Equilibration and Convergence of Lipid Bilayer Simulations . . . . . . . . . . S-12

S1.4 Form Factor . . . . . . . . . . . . . . . . . . . . . . . . . . . . . . . . . . . S-14

S1.5 `prosECCo75` Does Not Compromise the Original CHARMM Lipid Force Field Quality . . . . . . . . . . . . . . . . . . . . . . . . . . . . . . . . . . . . . . S-14

S1.5.1 POPC, POPE, POPS - Order Parameters, Form Factors, Area per Lipid S-14

S1.5.2 DPPC - Main Transition Temperature (TM) . . . . . . . . . . . . . . S-18

S1.5.3 Cholesterol - Order Parameters . . . . . . . . . . . . . . . . . . . . S-18

S1.5.4 Preferred Calcium Ion Binding Pattern and Lipid Complexation . . . S-20

S1.5.5 Calcium/Sodium Density Profiles . . . . . . . . . . . . . . . . . . . S-22

S1.5.6 POPC in `prosECCo75` is Structurally Better than `ECC-CHARMM36` . . . S-27

S1.5.7 Experimentally Measured Differences in Order Parameter Response to Calcium Ions between DPPC and POPC Membranes are not Reproduced by Force Fields . . . . . . . . . . . . . . . . . . . . . . . . . S-28

S1.5.8 Sodium Ions Overbind to POPS Membranes Even in `prosECCo75` Force Field - Potassium Comparison . . . . . . . . . . . . . . . . . . . . . . S-29

**S2 Supplementary Information Amino Acids and Proteins** **S-30**

S2.1 Osmotic Coefficient Calculations for Amino Acids and Small Peptides . . . . S-30

S2.2 prosECCo75 does not Compromise the Original CHARMM Protein Force Field Quality . . . S-32
S2.2.1 Additional Data Osmotic Coefficients, Amino Acids, Dipeptides, and Tripeptides . . . S-32
S2.2.2 Dipeptides - Ramachandran plot . . . S-34
S2.2.3 IDPs - Radius of Gyration and Coil Propensity . . . S-35
**S3 Supplementary Information Saccharides** **S-36**
S3.1 Simulated Saccharides . . . S-36
S3.2 Osmotic Coefficient Calculations and Experimental Results for Acidic Saccharides . . . S-36
S3.3 Experimental Determination Osmotic Coefficient for Acidic Saccharides . . . S-37
S3.4 prosECCo75 does not Compromise the Original CHARMM Saccharide Force Field Quality . . . S-38
S3.5 Data Osmotic Coefficient Calculation and Experiments Saccharides . . . S-39
**S4 Supplementary Information Ions** **S-40**
S4.1 Partial Charges for Ions . . . S-40
S4.2 Simulated Ions . . . S-41
S4.3 Comparison Neutron Scattering Data and Simulation for Ion Solutions . . . S-41
S4.4 Characterization "_s" Anion Series and K_s Cation . . . S-43
**References** **S-45**

### S1 Supplementary Information Membranes

#### S1.1 Partial Charges for Lipids

The partial charges developed for the PC, PE, and PS groups are presented in the schemata below.

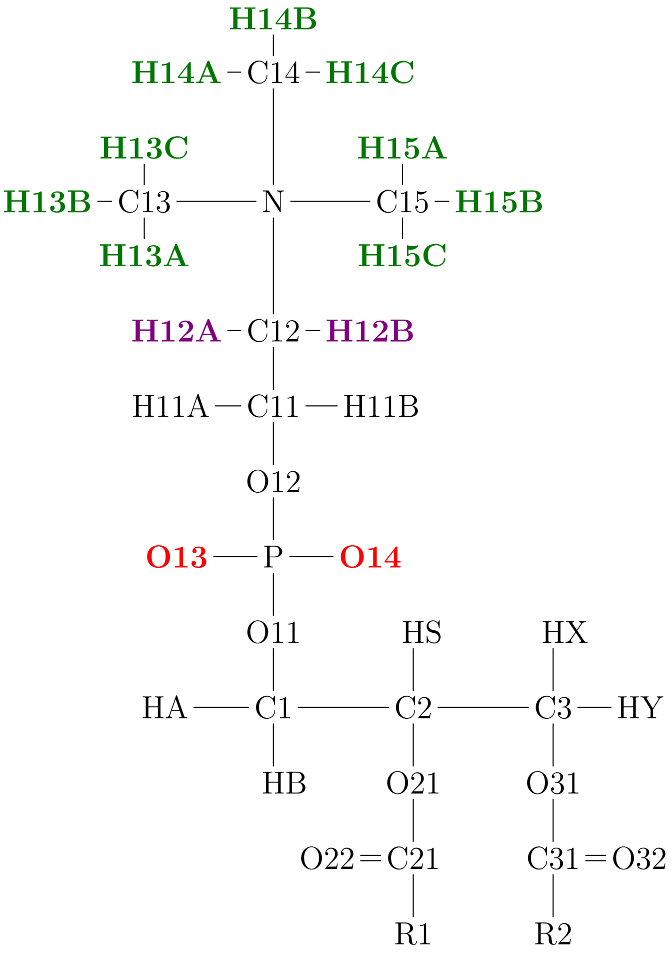


Figure S1: **PC.** The charge the oxygens was decreased by -0.125 from -0.780 to -0.655. The charges of this **hydrogens** were decreased by 0.02 from 0.25 to 0.23. The charges of this **hydrogens** were decreased by 0.035 from 0.25 to 0.215. These changes resulted in a total charge of +0.75 for the choline group and −0.75 for the phosphate group. The Lennard-Jones parameters were not changed.

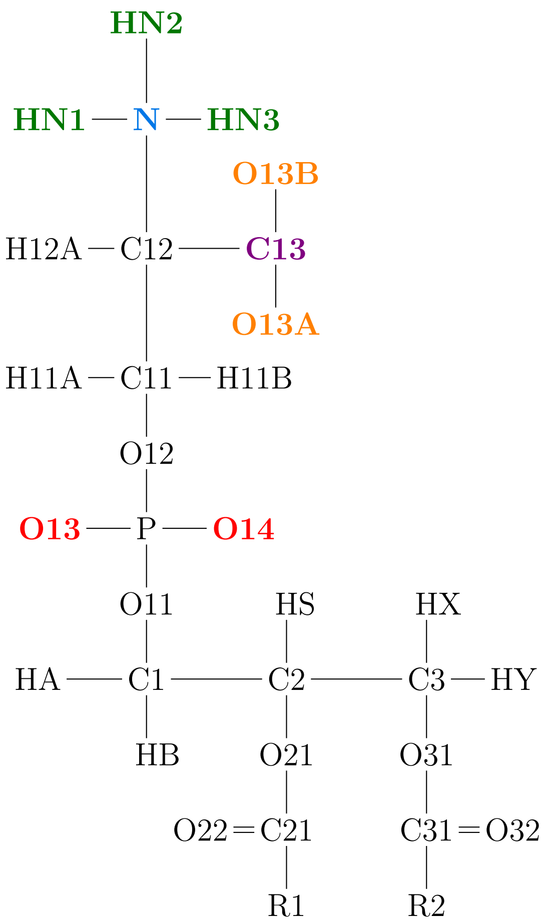


Figure S2: **PS.** The charge the **oxygens** was decreased by -0.125 from -0.780 to -0.655 . The charges of this **hydrogens** were decreased by 0.0825 from 0.33 to 0.2475. The charge of the **N** was increased from -0.3 to -0.3025. The charge on the **carbon** was increased by 0.05 from 0.34 to 0.39. The charges of **oxygens** were decreased by -0.1 from -0.67 to -0.57. These changes resulted in a total charge of +0.75 for the ammonium group, −0.75 for the carboxyl group, and −0.75 for the phosphate group. The Lennard-Jones parameters were not changed.

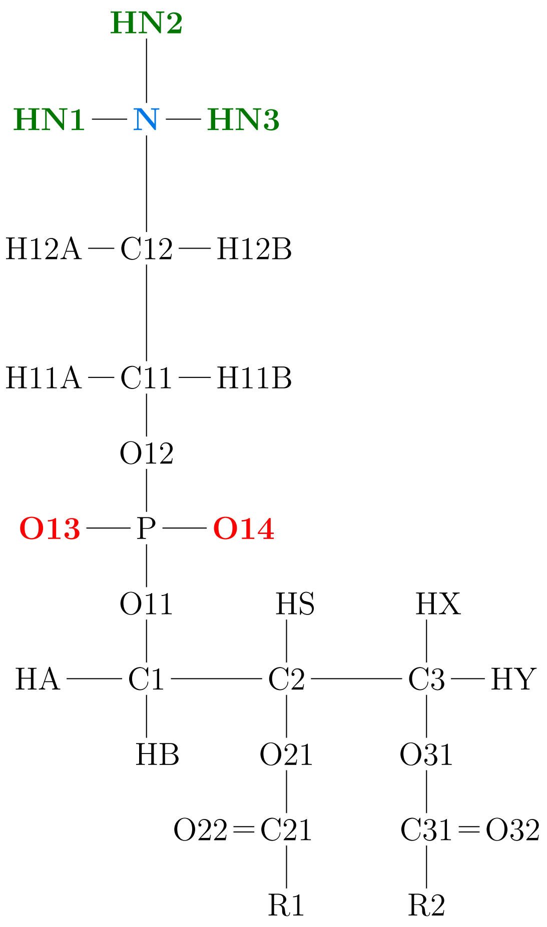


Figure S3: **PE.** The charge the **oxygens** was decreased by -0.125 from -0.780 to -0.655 . The charges of this **hydrogens** were decreased by 0.0825 from 0.33 to 0.2475. The charge of the **N** was increased from -0.3 to -0.3025. These changes resulted in a total charge of +0.75 for the ammonium group and −0.75 for the phosphate group. The Lennard-Jones parameters were not changed.

#### S1.2 Simulated Lipid Bilayer Systems

##### S1.2.1 POPC

Table S1: **Simulated POPC bilayers in different ion concentrations using the prosECCo75 force field** All systems contain 200 lipids and 9000 waters and were simulated at 310 K.

| **Salt** | **Total salt conc. [mM]** | **Bulk salt conc. [mM]** | **Num. of cations** | **Num. of anions** | **Time [ns]** | **Area per lipid[1] [nm$^{-2}$]** | **Data available** |
|---|---|---|---|---|---|---|---|
| None | 0 | 0 | 0 | 0 | 1370 | 0.627 | S1 |
| $CaCl_2$ | 25 | 12 | 4 | 8 | 1000 | 0.627 | S1 |
| $CaCl_2$ | 56 | 32 | 9 | 18 | 1000 | 0.621 | S1 |
| $CaCl_2$ | 111 | 80 | 18 | 36 | 1000 | 0.621 | S1 |
| $CaCl_2$ | 222 | 195 | 36 | 72 | 1000 | 0.615 | S1 |
| $CaCl_2$ | 450 | 449 | 73 | 146 | 1461 | 0.607 | S1 |
| $CaCl_2$ | 900 | 921 | 146 | 292 | 1000 | 0.595 | S1 |
| $CaCl_2$ | 2972 | 2997 | 496 | 992 | 1000 | 0.569 | S1 |
| NaCl | 56 | 44 | 9 | 9 | 432 | 0.623 | S1 |
| NaCl | 111 | 87 | 18 | 18 | 1000 | 0.603 | S2 |
| NaCl | 481 | 452 | 78 | 78 | 1000 | 0.627 | S1 |
| NaCl | 900 | 902 | 146 | 146 | 432 | 0.604 | S1 |
| NaCl | 968 | 985 | 157 | 157 | 174 | 0.595 | S1 |

[1] Standard error estimated using block averaging: $\pm 0.001\,\mathrm{nm}^{-2}$

Table S2: **Simulated POPC bilayers in different ion concentrations using the CHARMM36-NBFIX force field** All systems contained 200 lipids and 9000 water molecules and were simulated at 310 K.

| **Salt** | **Total salt conc. [mM]** | **Bulk salt conc. [mM]** | **Num. of cations** | **Num. of anions** | **Time [ns]** | **Area per lipid**[1] **[nm$^{-2}$]** | **Data available** |
|---|---|---|---|---|---|---|---|
| $CaCl_2$ | 56 | 35 | 9 | 18 | 595 | 0.648 | S3 |
| $CaCl_2$ | 111 | 94 | 18 | 36 | 1000 | 0.646 | S4 |
| $CaCl_2$ | 154 | 140 | 25 | 50 | 432 | 0.646 | S3 |
| $CaCl_2$ | 450 | 518 | 73 | 146 | 594 | 0.636 | S3 |
| NaCl | 56 | 42 | 9 | 9 | 667 | 0.650 | S3 |
| NaCl | 111 | 85 | 18 | 18 | 1000 | 0.628 | S5 |
| NaCl | 154 | 133 | 25 | 25 | 432 | 0.645 | S3 |
| NaCl | 968 | 1049 | 157 | 157 | 376 | 0.631 | S3 |

[1] Standard error estimated using block averaging: $\pm 0.001\,nm^{-2}$

Table S3: **Simulated POPC bilayers in different ion concentrations using the `CHARMM36 force field`** All systems contained 200 lipids and 9000 water molecules and were simulated at 310 K.

| **Salt** | **Total salt conc. [mM]** | **Bulk salt conc. [mM]** | **Num. of cations** | **Num. of anions** | **Time [ns]** | **Area per lipid**[1] **[nm$^{-2}$]** | **Data available** |
|---|---|---|---|---|---|---|---|
| None | 0 | 0 | 0 | 0 | 383 | 0.651 | S3 |
| $CaCl_2$ | 56 | 1 | 9 | 18 | 879 | 0.633 | S3 |
| $CaCl_2$ | 111 | 0 | 18 | 36 | 1000 | 0.619 | S6 |
| $CaCl_2$ | 450 | 56 | 73 | 146 | 897 | 0.582 | S3 |
| NaCl | 56 | 32 | 9 | 9 | 239 | 0.647 | S3 |
| NaCl | 111 | 70 | 18 | 18 | 1000 | 0.626 | S7 |
| NaCl | 968 | 998 | 157 | 157 | 863 | 0.629 | S3 |

[1] Standard error estimated using block averaging: $\pm 0.001\,nm^{-2}$

##### S1.2.2 DPPC

Table S4: **Data for DPPC bilayers without salt in different Temperatures using the `prosECCo75` and `CHARMM36` force field**

| System | Force Field | Total salt conc. [M] | Bulk salt conc. [M] | Num. of cations | Num. of anions | Time [ns] | Area per lipid[1] [$nm^{-2}$] | Temp. [K] | Available at |
|---|---|---|---|---|---|---|---|---|---|
| Pure | `prosECCo75` | 0 | 0 | 0 | 0 | 730 | 0.591 | 328[2] | S8 |
| $CaCl_2$ | `prosECCo75` | 0.450 | 0.430 | 73 | 146 | 860 | 0.479 | 328[2] | S9 |
| TM[2] | `prosECCo75` | 0 | 0 | 0 | 0 | 1000 | Fig ?? | 319-329[4] | S10 |
| TM[3] | `prosECCo75` | 0 | 0 | 0 | 0 | 1000 | Fig ?? | 319-329[4] | S11 |
| TM[2] | `CHARMM36` | 0 | 0 | 0 | 0 | 1000 | Fig ?? | 319-329[4] | S12 |
| TM[3] | `CHARMM36` | 0 | 0 | 0 | 0 | 1000 | Fig ?? | 319-329[4] | S13 |

[1] Standard error estimated using block averaging: $\pm 0.001\,nm^{-2}$
[2] Starting from a fluid conformation
[3] Starting from a gel conformation
[4] Every 1 K
[5] 128 DPPC molecules, 6400 water molecules

Table S5: **Data for DPPC bilayers with salt using the `ECC-DPPC` force field**. 128 lipid molecules were simulated in a box with 6400 water molecules. Run at 328 K

| System | Total salt conc. [M] | Bulk salt conc. [M] | Num. of cations | Num. of anions | Time [ns] |
|---|---|---|---|---|---|
| Pure | 0 | 0 | 0 | 0 | 300 |
| $CaCl_2$ | 0.160 | 0.120 | 19 | 38 | 570 |
| $CaCl_2$ | 0.380 | 0.290 | 39 | 78 | 1460 |
| $CaCl_2$ | 0.500 | 0.430 | 58 | 116 | 300 |
| $CaCl_2$ | 1.0000 | 0.860 | 116 | 232 | 300 |

##### S1.2.3 POPS and POPS/POPC Systems

The list of pure POPS and POPC:POPS 5:1 simulations that were performed for the structural validation (system with no ions), as well as the list of simulations concerning ion binding, is presented in Table S6.

Table S6: **Simulated POPS and POPC/POPS bilayers in different ion concentrations using the `prosECCo75` force field** All systems contains 144 lipids and 5760 waters and were simulated at 298 K.

| Salt | POPC | POPS | Contraions | Total salt conc. [mM] | Bulk salt conc. [mM] | Num. of cations[1] | Num. of anions | Time [ns] | Area per lipid[2] [$nm^{-2}$] | Available at |
|---|---|---|---|---|---|---|---|---|---|---|
| None | 0 | 144 | $Na^+$ 144 | 0 | 0 | 0 | 0 | 500 | 0.542 | S14 |
| None | 120 | 24 | $K^+$ 24 | 0 | 0 | 0 | 0 | 1200 | 0.610 | S15 |
| $CaCl_2$ | 120 | 24 | $K^+$ 24 | 96 | 24 | 10 | 20 | 1200 | 0.593 | S15 |
| $CaCl_2$ | 120 | 24 | $K^+$ 24 | 299 | 174 | 31 | 62 | 1200 | 0.580 | S15 |
| $CaCl_2$ | 120 | 24 | $K^+$ 24 | 1002 | 939 | 104 | 208 | 1200 | 0.565 | S15 |
| $CaCl_2$ | 120 | 24 | $K^+$ 24 | 2997 | 3090 | 311 | 622 | 1200 | 0.538 | S15 |
| None | 120 | 24 | $Na^+$ 24 | 0 | 0 | 0 | 0 | 500 | 0.561 | S16 |
| $CaCl_2$ | 120 | 24 | $Na^+$ 24 | 96 | 31 | 10 | 20 | 500 | 0.548 | S17 |
| $CaCl_2$ | 120 | 24 | $Na^+$ 24 | 299 | 212 | 31 | 62 | 500 | 0.539 | S18 |
| $CaCl_2$ | 120 | 24 | $Na^+$ 24 | 1002 | 982 | 104 | 208 | 500 | 0.527 | S19 |
| $CaCl_2$ | 120 | 24 | $Na^+$ 24 | 2997 | 3104 | 311 | 622 | 500 | 0.502 | S20 |

[1] In addition to counterions

[2] Standard error estimated using block averaging: $\pm 0.001\,nm^{-2}$

Table S7: **Simulated POPS and POPC/POPS bilayers in different ion concentrations using the `CHARMM36` force field** All systems contains 144 Lipids and 5760 water molecules and were simulated at 298 K. Unless stated otherwise.

| Salt | POPC | POPS | Contraions | Total salt conc. [mM] | Bulk salt conc. [mM] | Num. of cations[1] | Num. of anions | Time [ns] | Area per lipid[2] [$nm^{-2}$] | Available at |
|---|---|---|---|---|---|---|---|---|---|---|
| None | 0 | 128 | $Na^+$ 128 | 0 | 0 | 0 | 0 | 1000 | 0.546 | S21 |
| None | 24 | 120 | $Na^+$ 120 | 0 | 0 | 0 | 0 | 3000 | 0.575 | S22 |
| $CaCl_2$ | 24 | 120 | $Na^+$ 120 | 96 | 0.1 | 10 | 20 | 3000 | 0.556 | S23 |
| $CaCl_2$ | 24 | 120 | $Na^+$ 120 | 299 | 0.2 | 31 | 62 | 3000 | 0.529 | S24 |
| $CaCl_2$ | 24 | 120 | $Na^+$ 120 | 1002 | 336 | 104 | 208 | 3000 | 0.523 | S25 |
| $CaCl_2$ | 24 | 120 | $Na^+$ 120 | 2997 | 2566 | 311 | 622 | 3000 | 0.524 | S26 |

[1] In addition to counterions

[2] Standard error estimated using block averaging: $\pm 0.001\,nm^{-2}$

Table S8: **Simulated POPS and POPC/POPS bilayers in different ion concentrations using the `CHARMM36-NBFIX` (Han) force field.** All systems contain 5760 water molecules and were simulated at 298 K. Unless stated otherwise.

| Salt | POPC | POPS | Contraions | Total salt conc. [M] | Bulk salt conc. [M] | Num. of cations | Num. of anions | Time [ns] | Area per lipid[1] [$nm^{-2}$] | Available at |
|---|---|---|---|---|---|---|---|---|---|---|
| None | 250[2] | 50 | $Na^+$ 50 | 0 | 0 | 0 | 0 | 1200 | 0.649 | S27 |
| None | 120 | 24 | $Na^+$ 50 | 0 | 0 | 0 | 0 | 1000 | 0.581 | S28 |
| $CaCl_2$ | 120 | 24 | $Na^+$ 120 | 96 | 16 | 10 | 20 | 1000 | 0.576 | S29 |
| $CaCl_2$ | 250[3] | 50 | $Na^+$ 24 | 140 | 27 | 10 | 20 | 434 | 0.641 | S27 |
| $CaCl_2$ | 120 | 24 | $Na^+$ 120 | 299 | 178 | 31 | 62 | 1000 | 0.572 | S30 |
| $CaCl_2$ | 250[4] | 50 | $Na^+$ 24 | 940 | 946 | 31 | 62 | 437 | 0.628 | S27 |
| $CaCl_2$ | 120 | 24 | $Na^+$ 120 | 1002 | 1041 | 104 | 208 | 1000 | 0.556 | S31 |
| $CaCl_2$ | 120 | 24 | $Na^+$ 120 | 2997 | 3365 | 311 | 622 | 1000 | 0.520 | S32 |

[1] Standard error estimated using block averaging: $\pm 0.001\,nm^{-2}$
[2] 11207 water molecules, 320 K
[3] 11190 water molecules, 320 K
[4] 11174 water molecules, 320 K

##### S1.2.4 POPE Systems

Table S9: **Simulated POPE systems using the prosECCo75 force field** All the systems contain 144 lipids and 5760 water molecules and were simulated at 310 K.

| Salt | POPE | Contraions | Total salt conc. [M] | Bulk salt conc. [M] | Num. of cations | Num. of anions | Time [ns] | Area per lipid[1] [$nm^{-2}$] |
|---|---|---|---|---|---|---|---|---|
| None | 144 | None | 0 | 0 | 0 | 0 | 500 | 0.544 |

[1] Standard error estimated using block averaging: $\pm 0.001\,nm^{-2}$

##### S1.2.5 Systems with Cholesterol

Table S10: **Simulated POPC systems with cholesterol using the prosECCo75 force field** All the systems contain 200 POPC lipids and 5760 water molecules and were simulated at 310 K.

| Salt | Cholesterol | Contraions | Total salt conc. [M] | Bulk salt conc. [M] | Num. of cations | Num. of anions | Time [ns] | Area per lipid[1] [$nm^{-2}$] | Available at |
|---|---|---|---|---|---|---|---|---|---|
| None | 0 | None | 0 | 0 | 0 | 0 | 200 | 0.626 | S33 |
| None | 22 | None | 0 | 0 | 0 | 0 | 200 | 0.578 | S33 |
| None | 50 | None | 0 | 0 | 0 | 0 | 200 | 0.534 | S33 |
| None | 86 | None | 0 | 0 | 0 | 0 | 200 | 0.494 | S33 |
| None | 134 | None | 0 | 0 | 0 | 0 | 200 | 0.419 | S33 |
| None | 200 | None | 0 | 0 | 0 | 0 | 200 | 0.422 | S33 |

[1] Standard error estimated using block averaging: $\pm 0.001\,\mathrm{nm}^{-2}$

#### S1.3 Equilibration and Convergence of Lipid Bilayer Simulations

We checked the equilibration of the ion binding to POPC and POPC:POPS membranes. For `CHARMM36-NBFIX` and `prosECCo75`, the convergence is achieved within the first 100/200 ns for both cations, and show constant binding and unbinding events. For `CHARMM36` the situation is dramatically different with no unbinding event and much longer equilibration times, see Figures S4, S5, and S6.

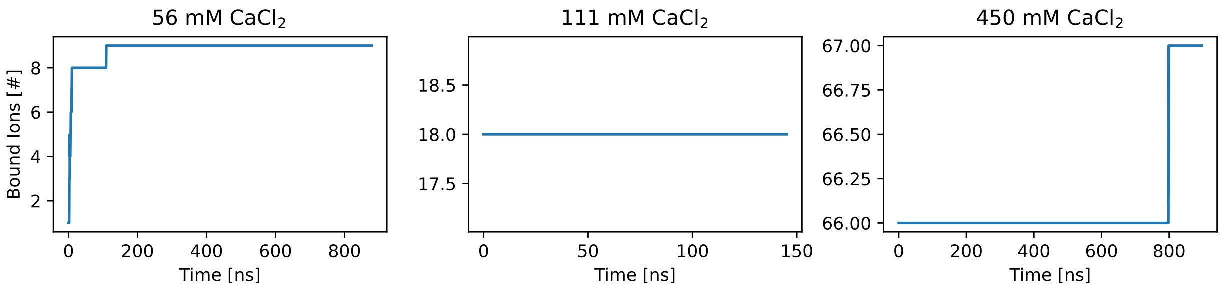


Figure S4: Binding of calcium ions to POPC membranes in `CHARMM36` force field as a function of time.

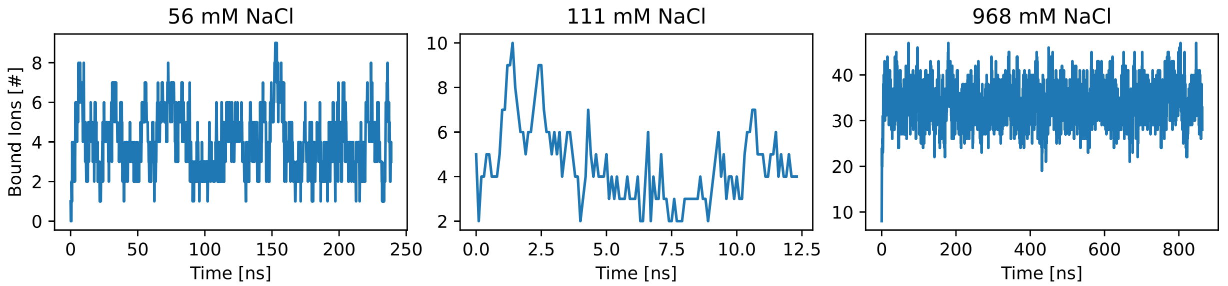


Figure S5: Binding of sodium ions to POPC membranes in CHARMM36 force field as a function of time.

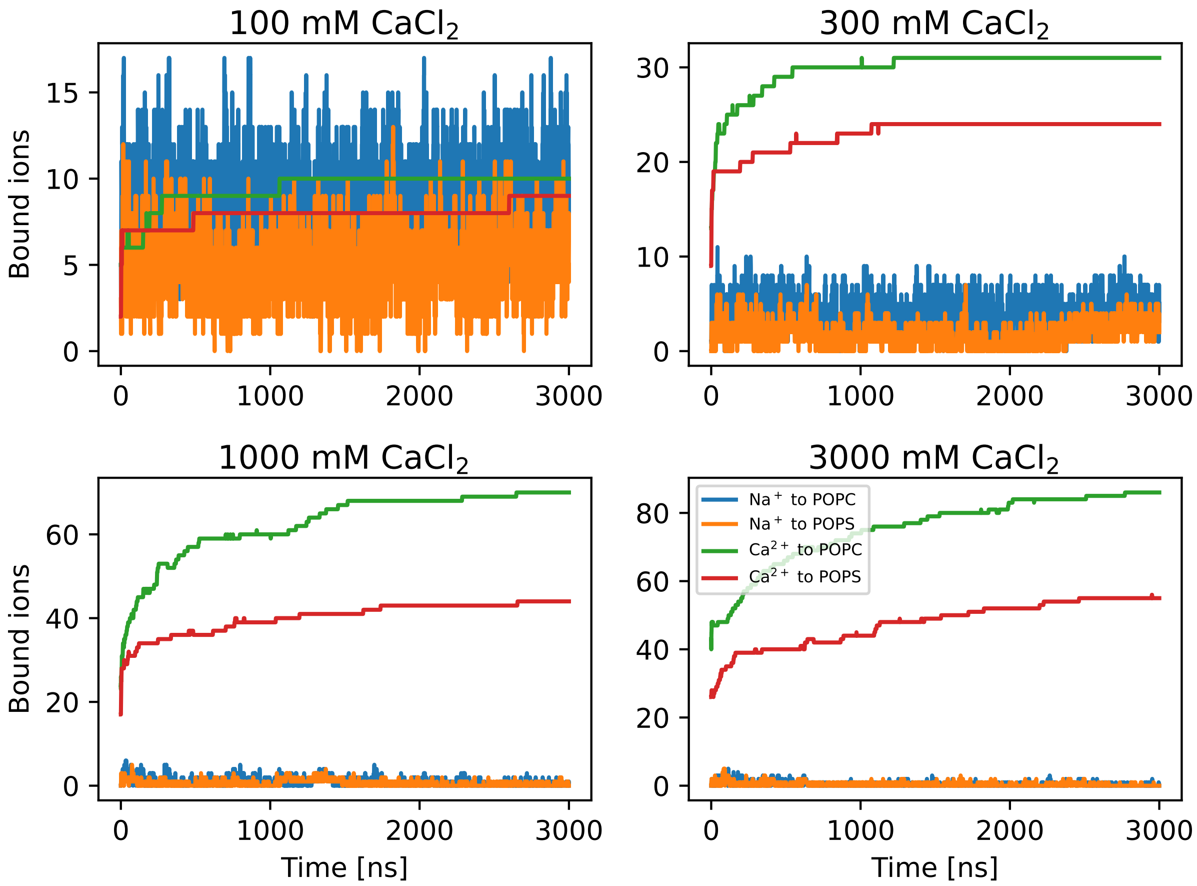


Figure S6: Binding of calcium ions and sodium counterions to 5:1 POPC:POPS membranes in CHARMM36 force field as a function of time. Ions are considered to be bound to POPC when they are closer than 0.325 nm of and POPC oxygen atom, correspondingly an ion is considered to be bound to POPS when it is closer than 0.325 nm of any POPS oxygen atom.

#### S1.4 Form Factor

To ensure that the structural properties of lipid membranes remain intact with `prosECCo75` implementation, we also examine scattering form factor profiles, $F(q)$, which offer information on the overall packing density and shape of the membrane. These form factors are calculated from simulations as Fourier transforms of electron density profiles, with the bulk value subtracted, $\rho(z) = \rho^{all}(z) - \rho^{bulk}$, along the z-direction

$$\parallel F(q) \parallel = \sqrt{\left(\sum_{i=0}^{N} \rho(z_i)(\cos(qz_i))\Delta z\right)^2 + \left(\sum_{i=0}^{N} \rho(z_i)(\sin(qz_i))\Delta z\right)^2}. \tag{S1}$$

The form factors obtained from X-ray and neutron scattering experiments are directly comparable with simulations. Additionally, scattering data can be analyzed using a model to derive parameters such as area per lipid and membrane thickness. However, these derived parameters are dependent on the chosen model, which may introduce uncertainties. Nevertheless, we provide a brief comparison between the simulation and experimental results in Table S11.

#### S1.5 `prosECCo75` Does Not Compromise the Original CHARMM Lipid Force Field Quality

##### S1.5.1 POPC, POPE, POPS - Order Parameters, Form Factors, Area per Lipid

With the `prosECCo75` approach, we aim to address a very specific overbinding issue by scaling the charges of the well-established CHARMM36 force field. While this approach benefits the extensive work done to fine-tune interactions in the original force field, our modifications may potentially disrupt this balance, potentially affecting `CHARMM36`'s excellent agreement with different experimental data.

To confirm that `prosECCo75` reproduces the experimental structural characteristics of lipid membranes with the same precision as `CHARMM36`, we calculated deuterium order parameters for lipid headgroups and acyl chains, along with form factors for the whole bilayers. These results were compared against experimental data, as shown in Figure S7. The headgroup order parameters describe the conformational space sampled of the headgroups, whereas the acyl chain order parameters describe the packing of lipid chains, and thus their fluidity. Form factors also reflect on the packing of the lipid chains. However, their direct interpretation is rather difficult, since they represent a Fourier transform of an electron density profile. Therefore we later compare also additional more intuitive structural parameters between CHARMM36 and `prosECCo75` and the experiment in Table S11.

As shown in the top left panel of Fig. S7, the scaling of charges in `prosECCo75` leaves the headgroup structure mainly unchanged. Slight improvements are observed for the $\beta$ and $g_1$ carbons, whereas $\gamma$ carbon is equally well presented by both models, and the $\alpha$, $g_3$, and $g_2$ carbons show a little worse agreement with experimental data with `prosECCo75`. It is likely caused by the decreased charge in the phosphate group, which affects the dihedral connecting the head group and the glycerol backbone. In the case of POPE and POPS lipids (top middle and right panels), some carbons are better described by `CHARMM36` and some by `prosECCo75`.

This indicates that such subtle changes might fall within the error estimates for the simulated and experimental data points and our `prosECCo75` implementation does not compromise the structure of headgroups of PC, PE and PS phospholipids while compared to the original `CHARMM36` model.

Similarly, the acyl chain order parameters do not significantly differ between our `prosECCo75` and `CHARMM36`, for none of the POPC, POPE and POPS lipid membranes, shown in the middle row of Fig. S7. Both `prosECCo75` and `CHARMM36` agree reasonably well with acyl chain order parameter experimental data for POPC and POPE lipid membranes. This means that the packing of the lipid bilayers is described reasonably in both models. This conclusion is also supported by a good agreement of both models with x-ray scattering form-factor experimental data. In the case of POPS membranes, both `prosECCo75` and `CHARMM36` models have

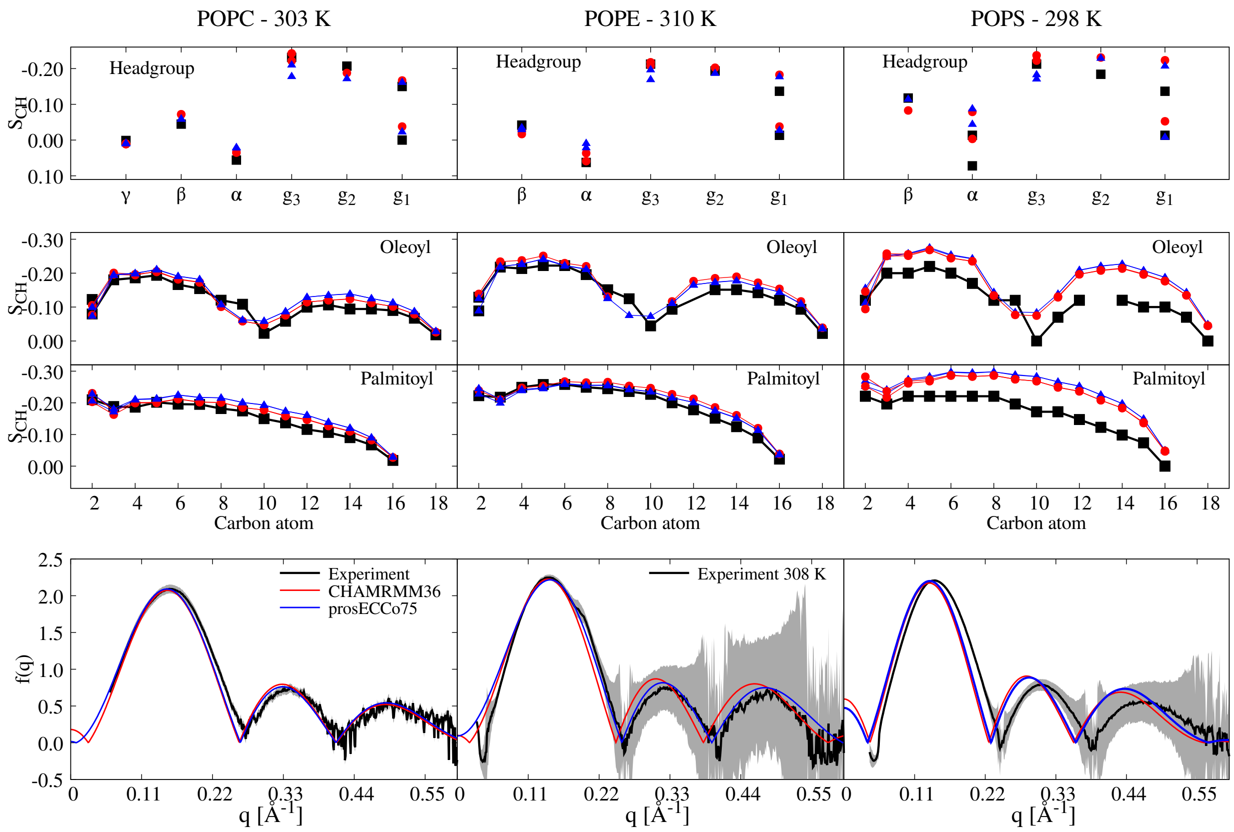


Figure S7: **Structural parameters from CHARMM36 and prosECCo75 models are compared with experimental values.** The top panels show headgroup and acyl chain order parameters. Experimental data for POPC are from Ref. S34. Experimental data for POPE headgroup order parameters are from Ref. S35. POPS experimental data for headgroup order parameters are from Ref. S21, for acyl chains from Ref. S36. POPS membranes are with sodium contraions. The bottom panel shows form factors. POPC CHARMM36 data are from Ref. S37 and experimental POPC data from Ref. S38. Experimental data for POPS are from Ref. S39 and experimental data for POPE from Ref. S40.

higher values of acyl chain order parameters than experimentally observed. This means that the membranes are overcondensated in both of these models. Again, this is in agreement with form-factor data where both simulation models have rather poor agreement with the experiment.

Table S11 lists the areas per lipid (APL) and thicknesses of DOPC, DPPC, POPS, and POPE membranes obtained from scattering experiments using a model and from MD simulations with the CHARMM36-NBFIX and prosECCo75 force fields.

Table S11: **Area per lipid and membrane thickness** Structural parameters remain close to the experimental range after ECC scaling.

| **Lipid** | **APL** `CHARMM36-NBFIX` **[nm$^2$]** | **APL** `prosECCo75` **[nm$^2$]** | **APL Experiment [nm$^2$]** | **Thickness** `CHARMM36-NBFIX` **[nm]** | **Thickness** `prosECCo75` **[nm]** | **Thickness Experiment [nm]** |
|---|---|---|---|---|---|---|
| POPC | 0.642 (303 K)<br>0.650 (310 K) | 0.619 (303 K)<br>0.627 (310 K) | 0.643 (303 K)[S38]<br>0.659 (310 K)[S38]<br>0.683 (303 K)[S41] | 3.88 (310 K) | 3.95 (303 K)<br>3.90 (310 K) | 3.91 (303 K)[S38] |
| DPPC | 0.609 (325 K) | 0.571 (323 K)<br>0.578 (324 K)<br>0.574 (325 K) | 0.635 (325 K)[S38]<br>0.628 (323 K)[S42] | 3.97 (325 K) | . | 3.9 (323 K)[S38] |
| POPS | 0.55 (298 K, $Na^+$)[S21]<br>0.56 (298 K, $K^+$)[S21] | 0.54 (298 K, $Na^+$)<br>0.60 (298 K, $K^+$) | 0.627 (298 K, $Na^+$)[S39] | 4.25 (298 K, $Na^+$)[S21]<br>4.25 (298 K, $K^+$)[S21] | 4.25 (298 K, $Na^+$)<br>3.95 (298 K, $K^+$) | 3.82 (298 K, $Na^+$)[S39] |
| POPE | 0.535 (310 K)[S43] | 0.544 (310 K) | 0.580 308 K[S40] | 3.52 (310 K)[S43] | 3.47 (310 K) | 3.99 (308 K)[S40] |

The areas per lipid of POPC are slightly smaller with `prosECCo75` than with `CHARMM36`, with the latter showing excellent agreement with some experimental values.[S38] In contrast, another experiment reported somewhat larger values.[S41] For DPPC, both `CHARMM36` and `prosECCo75` underestimate the APL values, in line with the behavior observed in earlier `CHARMM36` simulations performed with GROMACS near the main transition temperature of DPPC.[S44,S45] `prosECCo75` again predicts smaller APL values, likely due to the decreased inter-lipid repulsion caused by the scaled-down charges. For POPC and DPPC, the membrane thicknesses agree well between simulation models and experiments.

The APL values for POPS lipids are somewhat underestimated by both `CHARMM36` and `prosECCo75`. The fact that the latter shows improved agreement with the experiment when $K^+$ counter-ions are used instead of $Na^+$ ones indicates that the too-small APL values result from excessive condensation caused by the counter-ions. This hints that monovalent ions might bind too much to the PS headgroup not only in `CHARMM36` but also in `prosECCo75` despite charge-scaling. The too-small APL values of `CHARMM36` and `prosECCo75` for POPC go hand in hand with too-large membrane thickness values.

##### S1.5.2 DPPC - Main Transition Temperature (TM)

Phase behavior is a critical aspect of lipid bilayers. The main phase transition temperature ($T_\mathrm{m}$) is the temperature when a bilayer changes from the gel (*i.e.*, solid) to the liquid-disordered (*i.e.*, liquid) phase. To characterize how the changes in interaction parameters from `CHARMM36` to `prosECCo75` affect $T_\mathrm{m}$, we evaluated its value for a phosphatidylcholine lipid in `CHARMM36` (identical to `CHARMM36-NBFIX` in the absence of ions) and `prosECCo75`, and compared the obtained values to experiment. As POPC has a $T_\mathrm{m}$ (271 K) below the water freezing point, we use DPPC with a $T_\mathrm{m}$ of 314 K instead. Using two sets of simulations starting either from the liquid or the gel phase, respectively, we simulate a DPPC membrane at temperatures close to the experimental one and monitor the change of phase during the simulation. Clearly, both `CHARMM36/CHARMM36-NBFIX` ($T_\mathrm{m}$ between 319 and 325 K) and `prosECCo75` ($T_\mathrm{m}$ between 322 and 328 K) show very similar $T_\mathrm{m}$ values which are only slightly above the experimental value (see section S1.5.2 in the SI).

##### S1.5.3 Cholesterol - Order Parameters

In addition to PC, PE, and PS that form the majority of the phospholipids in the plasma membrane and other cellular membranes,[S46] cholesterol is also present in concentrations between 10% and 50% in these membranes.[S46] Therefore, the accurate modeling of phospholipid–cholesterol interactions is also crucial for a lipid force field. We evaluated the effect of cholesterol on the order parameters of POPC headgroups and acyl chains in both CHARMM36 and `prosECCo75` force fields and compared them to the experiment. As demonstrated previously,[S34,S47] cholesterol has little effect on the headgroup conformations in experiments, and this behavior is relatively well reproduced by the CHARMM36 force field.[S48] The data, shown in Fig. S9, demonstrates that the scaling of the PC headgroup charges based on the ECC approach has little effect on the headgroup order parameters in a POPC/cholesterol 50/50 mixture. The order parameters for the $g_3$ carbon deviate slightly between the two models. Yet, this behavior is likely independent of cholesterol, as it was already observed for

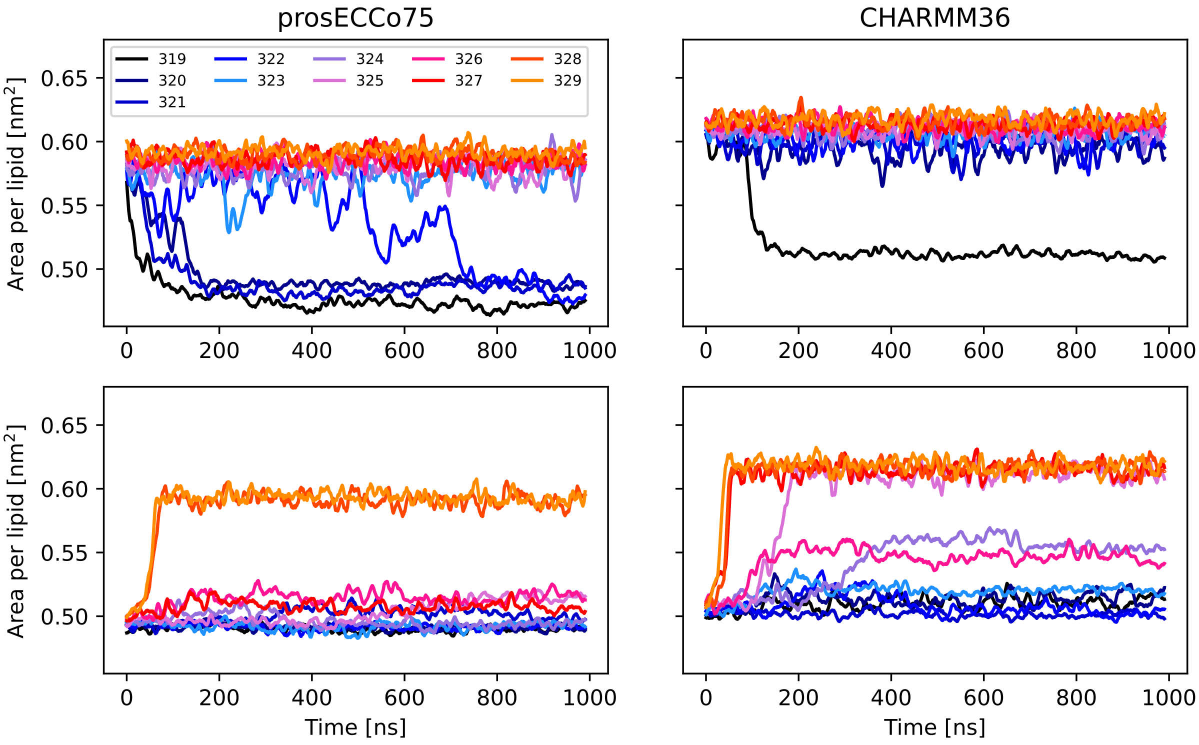


Figure S8: Simulations of DPPC at different temperatures using `prosECCo75` and `CHARMM36` stating from the liquid phase and the gel phase to determine the main phase transition temperature. Running average over 10 ns is shown.

pure POPC bilayers (see Fig. S7). These results highlight that the scaling of the headgroup charges does not compromise the lipid–cholesterol interactions in the `prosECCo75` force field, and it can thus be safely used to model lipid bilayers containing cholesterol.

Overall, the `prosECCo75` and `CHARMM36/CHARMM36-NBFIX` force fields reproduce the experimental properties of all studied lipid bilayers with comparable accuracy without salt. However, as demonstrated in the main text, `prosECCo75` also reproduces experimental behavior in their presence, thus rendering it a better choice to model biomembranes in their native conditions.

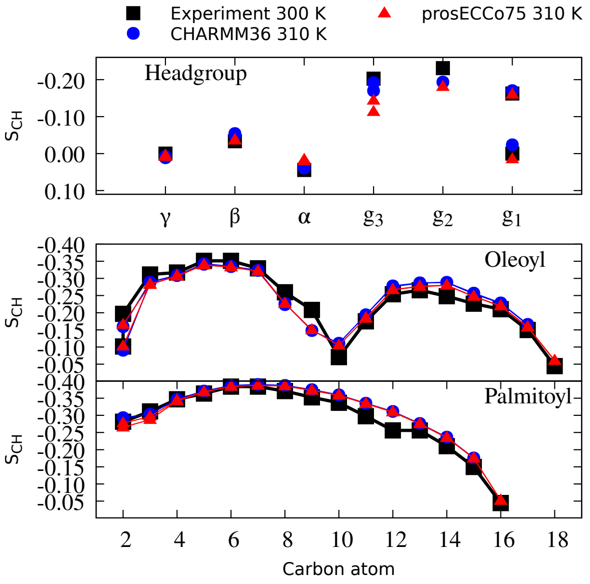


Figure S9: **Order Parameters of POPC molecule in POPC/cholesterol 1:1 mixture.** Both the original CHARMM36 and its scaled version prosECCo75 reproduce the experimental data with similar accuracy except for the g3 carbon–hydrogen bond order parameter. Here, CHARMM36 performs better.

##### S1.5.4 Preferred Calcium Ion Binding Pattern and Lipid Complexation

The complexation number of an ion is calculated as the number of lipid phosphate atoms located within a distance of 0.5 nm from the specified ion. The cut-off value was previously validated.[S49]

Although the response of the headgroup order parameters to calcium ions is similar in both the `CHARMM36-NBFIX` and `prosECCo75` force fields, there are clear differences between these force fields in terms of coordination numbers and residency times. In the `prosECCo75` force field, the mean coordination number is ≈ 1.7, which is higher than that of `CHARMM36-NBFIX` with a mean coordination number of 1.3. Both values are somewhat lower than 2, which was obtained from fitting binding models to experiments.[S50] In contrast, `CHARMM36` has an average coordination number of 3.5, significantly exceeding the experimental estimate.

The interaction of $Ca^{2+}$ with different oxygen atoms also varies significantly among the

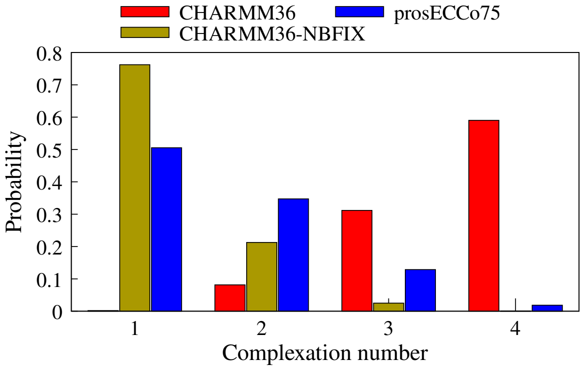


Figure S10: **Complexation of POPC lipids to calcium ions.** The coordination number of calcium ions with POPC molecules was determined as the number of lipids in which phosphate or carbonyl oxygen atoms are within the cut-off of 0.5 nm of a given ion. Reported systems have 450 mM calcium system concentration.

models, as illustrated in Table S12. In `CHARMM36`, $Ca^{2+}$ primarily interacts with phosphate oxygens without penetrating deeper into the carbonyl region. In CHARMM36-NBFIX, cations exhibit less tight binding to phosphates, allowing them to transiently sample the carbonyl region deeper within the membrane. In `prosECCo75`, $Ca^{2+}$ samples the carbonyl region even more effectively.

Table S12: **Ion binding to oxygen atoms of phosphate and carbonyl group of POPC lipids**. Top: Sodium ions binding to POPC membrane with 968 mM system concentration of NaCl. Bottom: Calcium ion binding to POPC membrane with 450 mM system concentration of $CaCl_2$. The cutoff used is 0.325 nm.[S49]

| $Na^+$ | Phosphate oxygens | Carbonyl oxygens |
|---|---|---|
| CHARMM36 | 81 % | 52 % |
| CHARMM36-NBFIX | 61 % | 62 % |
| prosECCo75 | 81 % | 63 % |

| $Ca^{2+}$ | Phosphate oxygens | Carbonyl oxygens |
|---|---|---|
| CHARMM36 | 100 % | 3 % |
| CHARM36-NBFIX | 88 % | 24 % |
| prosECCo75 | 90 % | 52 % |

##### S1.5.5 Calcium/Sodium Density Profiles

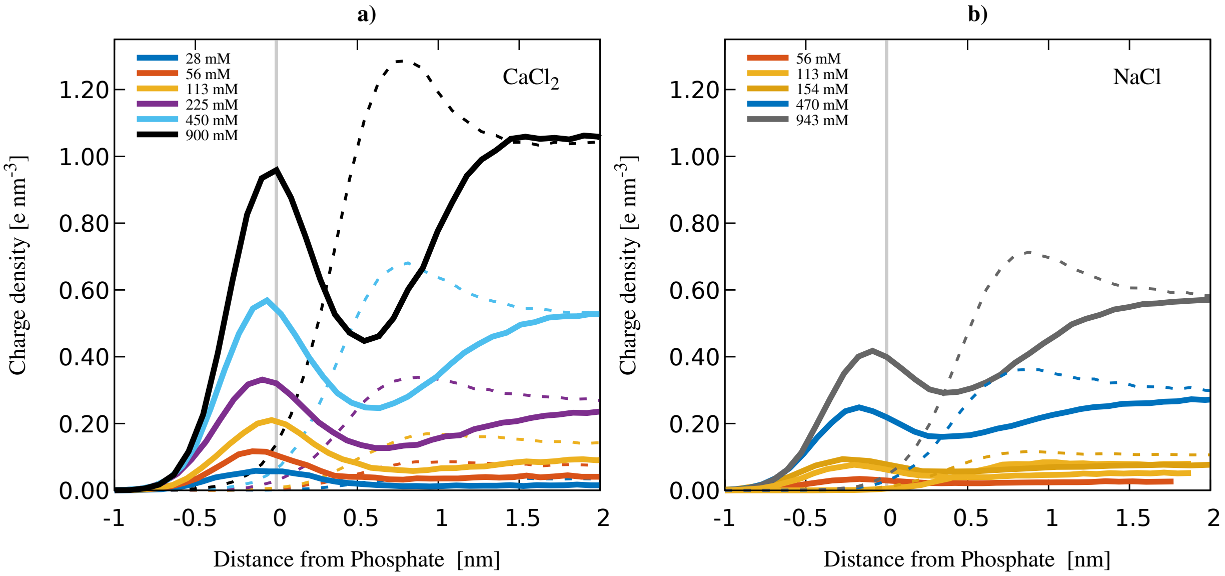


Figure S11: Density profiles of calcium and sodium ion binding to POPC membranes at different concentrations run with prosECCo75 force field.

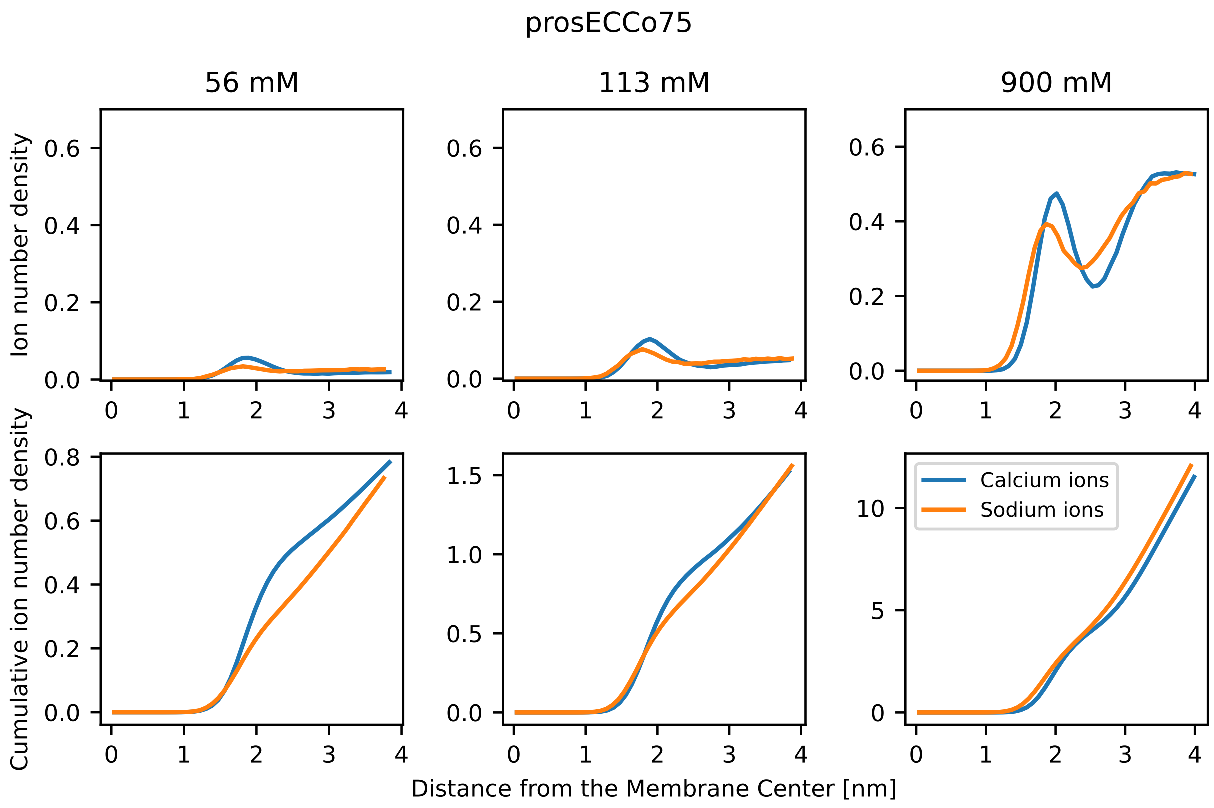


Figure S12: Density profiles of calcium and sodium ion binding to POPC membranes at different concentrations run with `prosECCo75` force field. Top panels: Number density profiles of ions along the membrane normal. The membrane center is at 0 nm. Bottom panels: Integrated ion number density along the membrane normal. Concentrations reported refer to the system concentration.

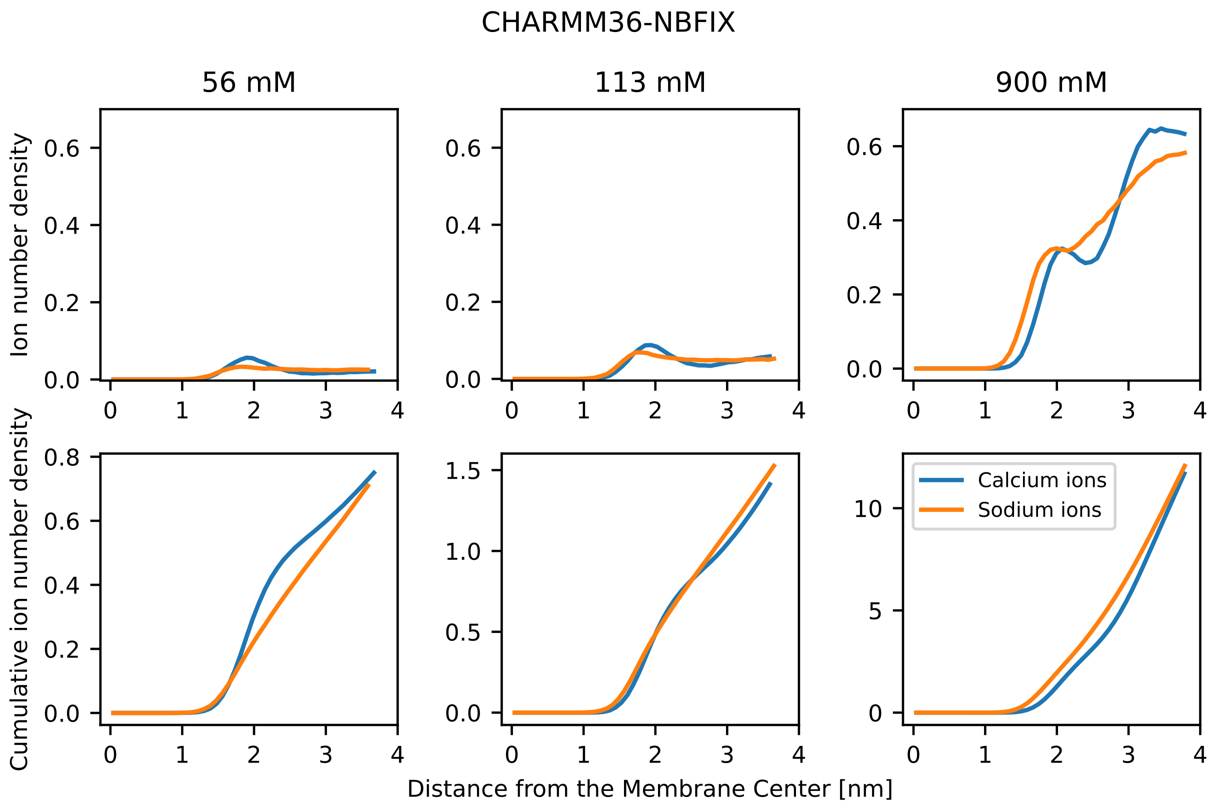


Figure S13: Density profiles of calcium and sodium ion binding to POPC membranes at different concentrations run with `CHARMM36-NBFIX` force field. Top panels: Number density profiles of ions along the membrane normal. The membrane center is at 0 nm. Bottom panels: Integrated ion number density along the membrane normal. Concentrations reported refer to the system concentration.

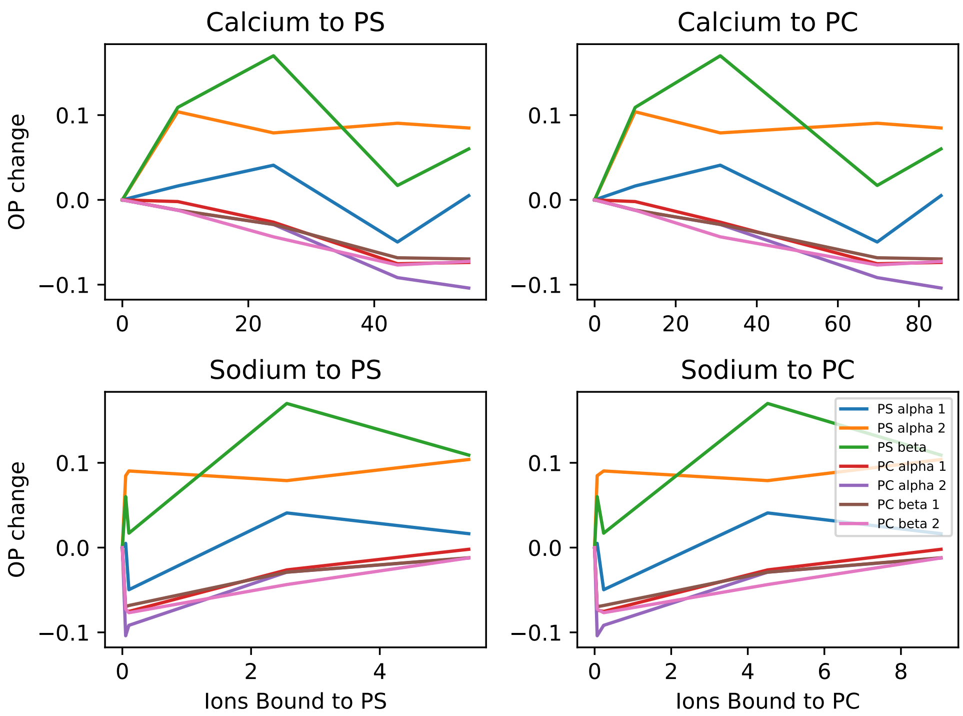


Figure S14: POPS Headgroup order parameter change with ions bound to the membrane is not trivial in `CHARMM36` model of POPC:POPS membranes.

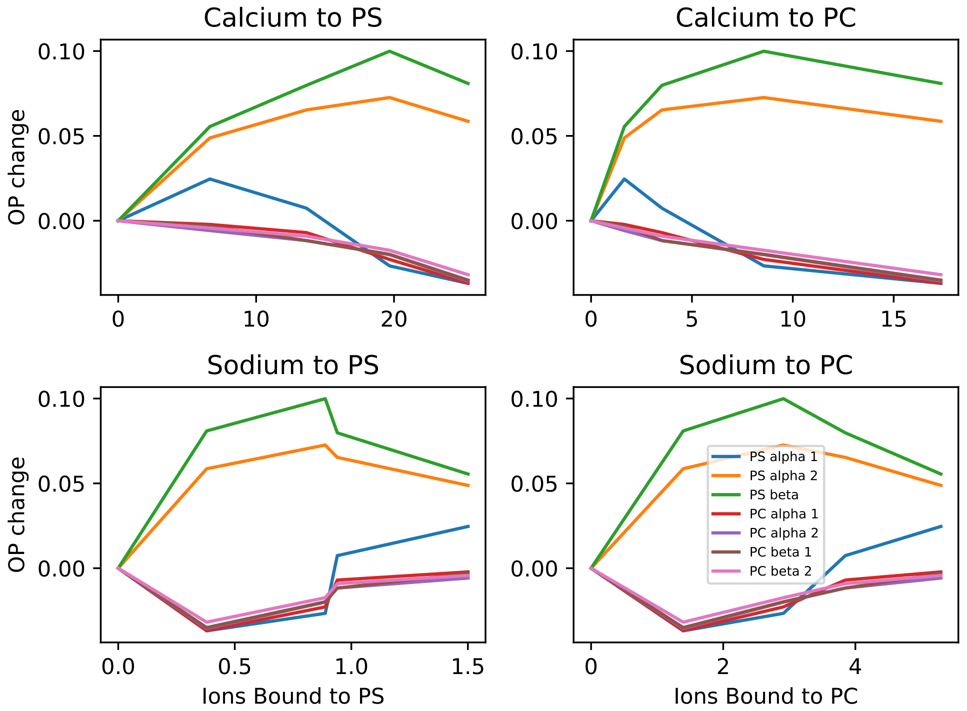


Figure S15: POPS Headgroup order parameter change with ions bound to the membrane is not trivial in CHARMM36-NBFIX model of POPC:POPS membranes.

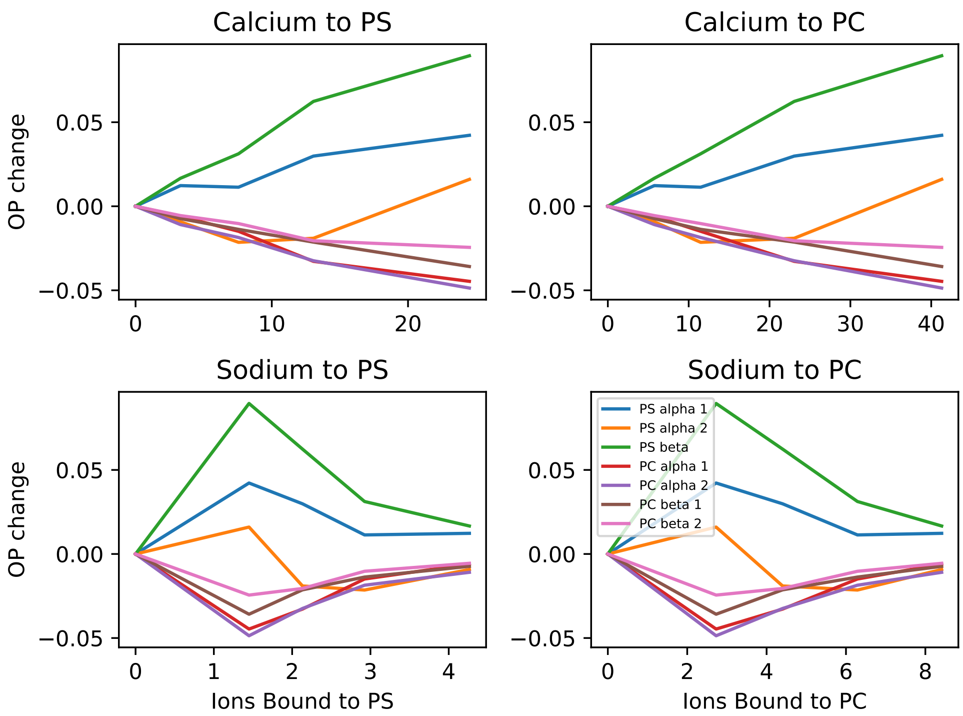


Figure S16: POPS Headgroup order parameter change with ions bound to the membrane is not trivial in `prosECCo75` model of POPC:POPS membranes.

##### S1.5.6 POPC in `prosECCo75` is Structurally Better than `ECC-CHARMM36`

Our previous model of POPC using ECC, named `ECC-CHARMM36`,[S51] followed the recipe optimized for amber14 POPC force field.[S51] Because it scaled every partial charge and Lennard-Jones parameters, several dihedrals were affected, see Figure S17. `prosECCo75` behaves better as we purposely do not touch so many dihedrals, especially critical ones.

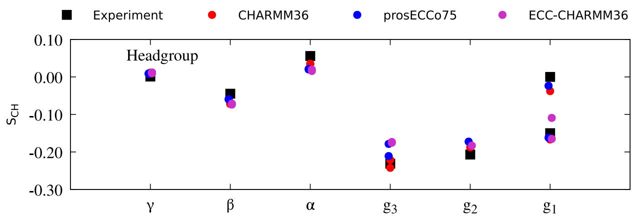


Figure S17

##### S1.5.7 Experimentally Measured Differences in Order Parameter Response to Calcium Ions between DPPC and POPC Membranes are not Reproduced by Force Fields

Interestingly, the effect of calcium ions concentration on the experimental NMR $\alpha$-CH bond order parameter differs slightly between DPPC[S52] and POPC[S50] membranes with DPPC headgroups being more sensitive to calcium ions. This difference between the response of $\alpha$ order parameter to calcium ions in DPPC (see Table S5 and POPC membranes is not reproduced by the `prosECCo75` model nor by ECC-POPC amber-based force-field,[S51] see Figure S18.

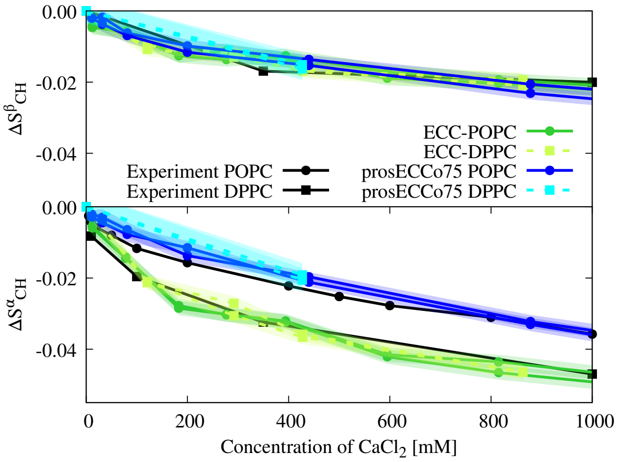


Figure S18: $\alpha$-CH and $\beta$-CH response to calcium ions in POPC and DPPC membranes. No difference in the behavior is observed between POPC and DPPC membranes in simulations. Both CHARMM36-based prosECCo75 force field and Amber-based ECC-POPC, ECC-DPPC force field were examined. Experimental data for DPPC was measured at 323 K and taken from DPPC.[S52] Experimental data for POPC was measured at 313 K and taken from.[S50] DPPC simulations were run at 328 K. POPC simulations were run at 310 K.

While both POPC and DPPC simulations with `prosECCo75` force field follow the POPC experimental line, in the case of ECC-POPC forcefield, both POPC and DPPC simulations follow the DPPC experimental line.

In both force fields, `prosECCo75` and ECC-POPC, $\beta$-CH order parameter response to calcium ions is the same for POPC and DPPC membranes and follows the DPPC experimental line. Since no experimental data exist for the $\beta$-CH order parameter response to calcium ions for POPC membranes, it remains unclear if an experimental difference between the DPPC and POPC response to calcium ions exists also for $\beta$-CH order parameter.

##### S1.5.8 Sodium Ions Overbind to POPS Membranes Even in `prosECCo75` Force Field - Potassium Comparison

In `prosECCo75`, sodium counterions bind to pure POPS membranes significantly more than potassium counterions as can be seen in Figure S19 b). At the same time, the membranes simulated with potassium counterions have significantly better agreement of scattering form factors with experimental data, as in Figure S19 a). This indicates that sodium binding to pure POPS membrane is still overestimated in the prosECCo75 force field.

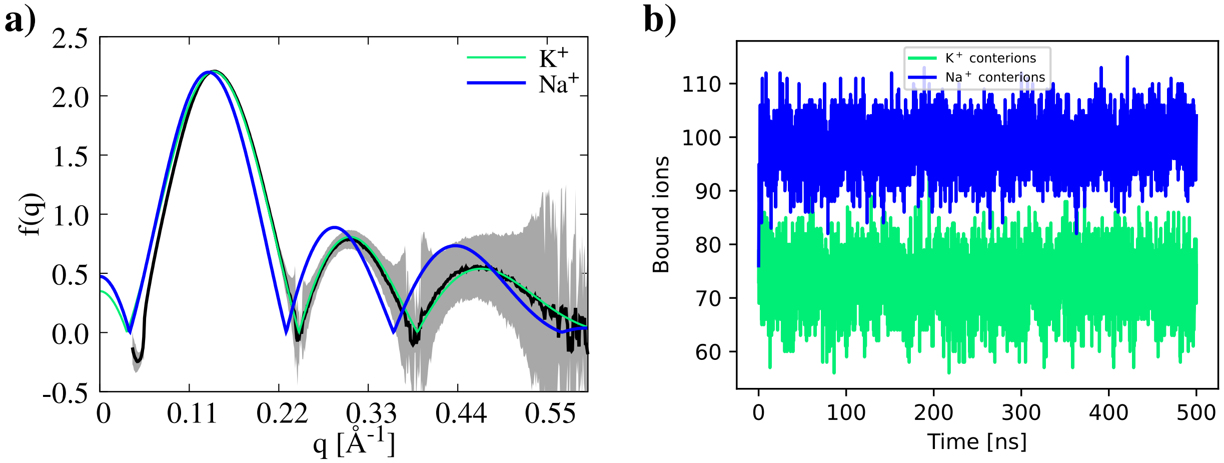


Figure S19: **POPS membrane with K$^+$ and Na$^+$ contraions simulated with `prosECCo75` force field a)** Form factor of POPS membrane with different contraions. **b)** Binding of different counterions to POPS membrane in time. Simulations are at 298 K.

### S2 Supplementary Information Amino Acids and Proteins

#### S2.1 Osmotic Coefficient Calculations for Amino Acids and Small Peptides

Osmotic coefficient calculations were employed to evaluate the performance of `prosECCo75` force field in modeling small organic molecules. We carried out simulations of twenty natural L-amino acids. Three concentrations were simulated for amino acid solutions [0.5 M; 1 M; 2M], similar to the previous study.[S53] We also adopted the same experimental reference data. In the case of charged species, the system was charge-neutralized by counter ions (sodium or chloride).

To calculate the osmotic coefficients, we adopted the methodology developed by Luo and Roux[S54] and further applied to amino acids and saccharides.[S53,S55] The solutes were randomly placed in a cubic box of 4.8 nm. Then, the simulation box was extended by 2.4 nm in each direction along the z-axis and filled with pure water. The solutes were restrained to the central 4.8 nm cube by a flat-bottom harmonic potential of $k = 1000$ kJ/mol$^{-1}$nm$^{-2}$. These virtual walls were acting only on solute atoms, while water molecules were free to move. Simulations were performed in NPT ensemble with the pressure coupling applied only to $z$-axis. The applied forces were extracted from the production MD part using the following equation:

$$F = \sum_{i}^{N} 1/2 \times k \times |z_i - z_{\text{wall}}| \quad \text{(S2)}$$

where $z_i$ is the instantaneous coordinate of atom $i$ and $z_{wall}$ is the position of the corresponding harmonic restraint. The summation runs over all the atoms $N$ that crossed the walls. The resulting osmotic pressure averaged over two walls was calculated as:

$$\Pi = 1/2 \times \langle F \rangle / A \quad \text{(S3)}$$

where $\langle F \rangle$ is the mean force over all analyzed frames, and $A$ is the cross-sectional area of the system.

Finally, molal osmotic coefficients were calculated using the ideal osmotic pressure ($\Pi_{\mathrm{id}}$) according to the equation

$$\phi = \frac{\Pi}{\Pi_{\mathrm{id}}} = \frac{\Pi}{RT\, m\nu\, \frac{M_w}{V_w}}, \tag{S4}$$

where $V_w$ is the molar volume of water (0.018 L·mol$^{-1}$), $R$ is the universal gas constant, $T$ is the absolute temperature, $m$ is the molality of the solute in the restrained part of the box, $\nu$ is the Van't Hoff coefficient (1 for neutral species and 2 for charged ones), and $M_w$ is the molar mass of water, 0.018 kg·mol$^{-1}$. Solute molality was calculated from the known number of solute molecules and by counting water oxygen atoms in the restrained region. Standard error was estimated using block averaging.

#### S2.2 prosECCo75 does not Compromise the Original CHARMM Protein Force Field Quality

##### S2.2.1 Additional Data Osmotic Coefficients, Amino Acids, Dipeptides, and Tripeptides

Table S13: Amino acids osmotic coefficients. *c36* corresponds to CHARMM36m, *p75* corresponds to prosECCo75, *exp* corresponds to experiments.[S53,S56]

| Concentration | 0.5 M | | | 1.0 M | | | 2.0 M | | |
|---|---|---|---|---|---|---|---|---|---|
| Amino Acid | *c36* | *p75* | *exp* | *c36* | *p75* | *exp* | *c36* | *p75* | *exp* |
| Ala (A) | 0.837 | 1.023 | 1.004 | 0.748 | 1.057 | 1.009 | 0.578 | 1.124 | 1.016 |
| Asn (N) | 0.874 | 1.014 | | 0.748 | 1.054 | | 0.543 | 1.002 | |
| Cys (C) | 0.817 | 0.994 | 0.886 | 0.706 | 0.981 | 0.736 | 0.518 | 0.964 | |
| Gln (Q) | 0.928 | 1.143 | | 0.751 | 1.057 | | 0.673 | 1.092 | |
| Gly (G) | 0.780 | 1.047 | 0.957 | 0.624 | 1.032 | 0.929 | 0.450 | 1.025 | 0.904 |
| His (H) | 0.951 | 1.179 | | 0.891 | 1.050 | | 0.794 | 1.123 | |
| Ile (I) | 0.881 | 1.091 | | 0.830 | 1.066 | | 0.578 | 1.075 | |
| Met (M) | 0.859 | 1.031 | | 0.655 | 0.959 | | 0.270 | 0.807 | |
| Leu (L) | 0.890 | 1.078 | | 0.842 | 1.058 | | 0.500 | 0.988 | |
| Phe (F) | 0.912 | 1.117 | | 0.713 | 0.936 | | 0.418 | 0.765 | |
| Pro (P) | 1.101 | 1.107 | 1.023 | 1.140 | 1.182 | 1.046 | 1.232 | 1.332 | 1.095 |
| Ser (S) | 0.852 | 1.044 | 0.950 | 0.691 | 1.018 | 0.907 | 0.456 | 0.994 | |
| Thr (T) | 0.918 | 1.063 | 0.988 | 0.781 | 1.011 | 0.982 | 0.586 | 1.060 | 0.979 |
| Trp (W) | 0.783 | 0.792 | | 0.496 | 0.608 | | 0.314 | 0.442 | |
| Tyr (Y) | 0.852 | 0.912 | | 0.653 | 0.780 | | 0.454 | 0.703 | |
| Val (V) | 0.980 | 1.100 | 1.036 | 0.869 | 1.089 | 1.073 | 0.735 | 1.159 | 1.146 |
| Arg (R) | 0.759 | 1.015 | 0.913 | 0.573 | 0.949 | 0.867 | 0.539 | 0.955 | |
| Glu (E) | 0.840 | 0.955 | 0.907 | 0.793 | 0.912 | 0.920 | 0.786 | 0.858 | 0.992 |
| Lys (K) | 0.790 | 1.046 | 0.917 | 0.746 | 1.104 | 0.980 | 0.784 | 1.275 | 1.130 |
| Asp (D) | 0.847 | 0.938 | | 0.701 | 0.886 | | 0.692 | 0.780 | |

Table S14: Dipeptide osmotic coefficients. *c36* corresponds to `CHARMM36-NBFIX`, *p75* corresponds to `prosECCo75`, *exp* corresponds to `experiments`.[S53,S56]

| Concentration | 1 M | | |
|---|---|---|---|
| dipeptide | *c36* | *p75* | *exp* |
| AA | 0.6662 | 1.1265 | 1.04 |
| GG | 0.2854 | 0.9283 | 0.882 |
| AG | 0.5665 | 1.035 | 0.956 |
| GA | 0.4173 | 1.0456 | 0.951 |

Table S15: Tripeptide osmotic coefficients. *c36* corresponds to `CHARMM36-NBFIX`, *p75* corresponds to `prosECCo75`, *exp* corresponds to `experiments`.[S53,S56]

| Concentration | 0.3 M | | |
|---|---|---|---|
| tripeptide | *c36* | *p75* | *exp* |
| GGG | 0.6815 | 1.0663 | 0.905 |

##### S2.2.2 Dipeptides - Ramachandran plot

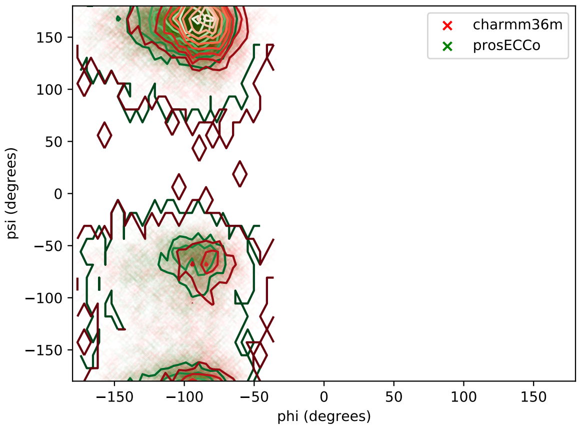


(a) RR

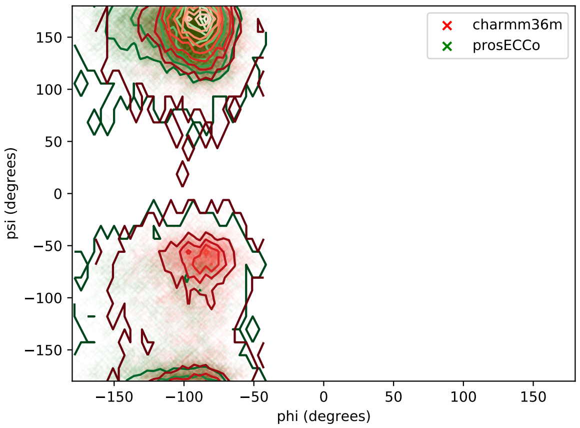


(b) KK

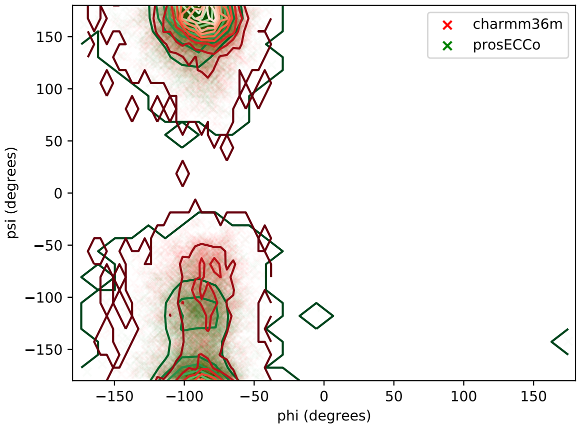


(c) DD

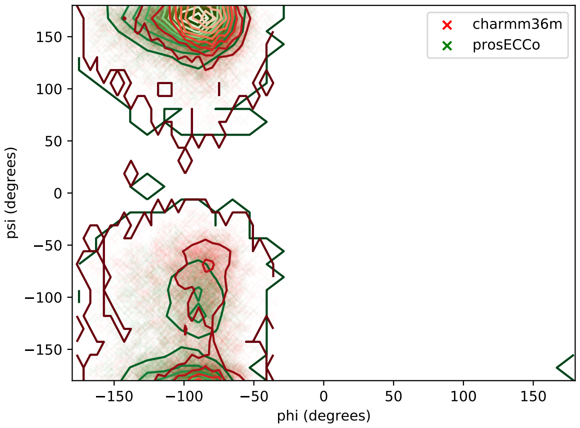


(d) EE

Figure S20: Ramachandran plot for four dipeptides: RR (a), KK(b), DD (c), and EE (d). The Ramachandran plot was calculated for the original FF (`CHARMM36-NBFIX`, red) and the scaled one (`prosECCo75`, green).

##### S2.2.3 IDPs - Radius of Gyration and Coil Propensity

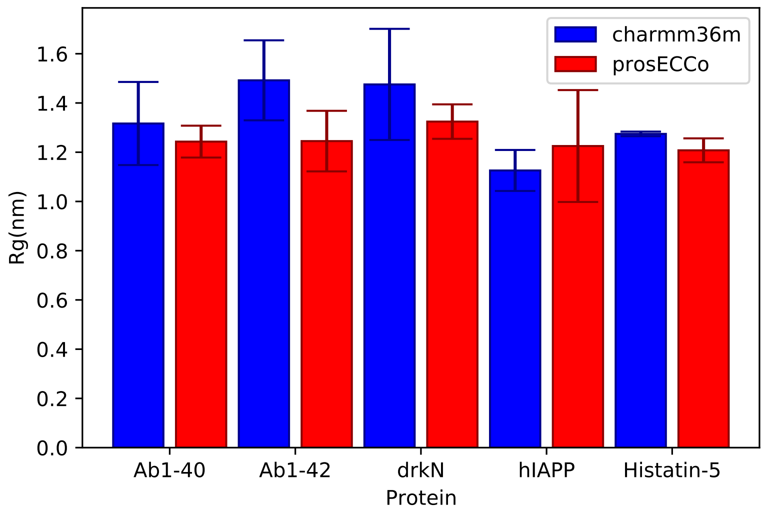


Figure S21: Gyration Radii for 5 IDPs: Abeta1-40, Abeta1-42, drkN, hIAPP and Histadin-5. The gyration radius was calculated with the original FF (`CHARMM36-NBFIX`, blue) and with the scaled one (`prosECCo75`, red).

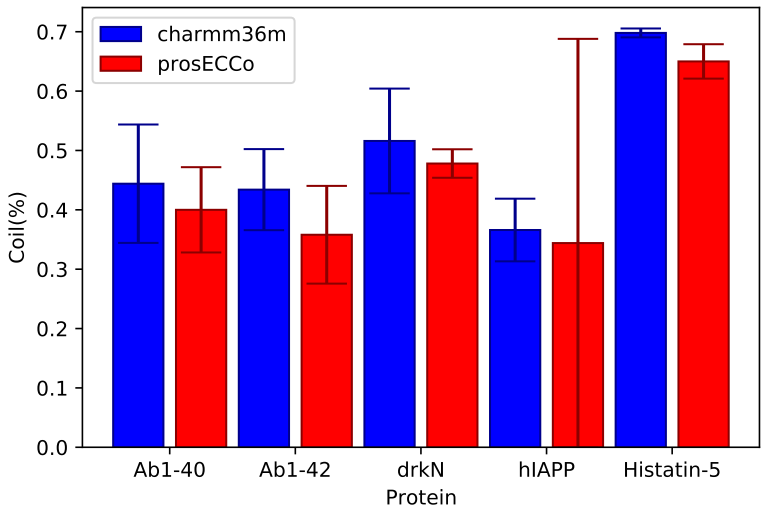


Figure S22: Percentage of random coiled structure for 5 IDPs: Abeta1-40, Abeta1-42, drkN, hIAPP, and Histadin-5. The coiled percentage was calculated with the original FF (`CHARMM36-NBFIX`, blue) and with the scaled one (`prosECCo75`, red).

### S3 Supplementary Information Saccharides

#### S3.1 Simulated Saccharides

Table S16: **Simulated carbohydrate solutions for osmotic coefficient predictions** The temperature in all simulations was 310 K.

| Carbohydrate | Designed molarity [M] | Num. of $\alpha$-isomers[1] | Num. of $\beta$-isomers[1] | Num. of $Na^+$ ions | Num. of waters | Time[2] [ns] | Zenodo link |
|---|---|---|---|---|---|---|---|
| D-GalA | 0.10 | 3 | 4 | 7 | 7197 | >200 | 10.5281/zenodo.10151774 |
| D-GalA | 0.25 | 6 | 10 | 16 | 7103 | >200 | 10.5281/zenodo.10151774 |
| D-GalA | 0.50 | 13 | 20 | 33 | 6936 | >200 | 10.5281/zenodo.10151774 |
| D-GalA | 0.75 | 19 | 31 | 50 | 6778 | >200 | 10.5281/zenodo.10151774 |
| D-GalA | 1.00 | 26 | 41 | 67 | 6614 | >200 | 10.5281/zenodo.10151774 |
| D-GlcA | 0.10 | 3 | 4 | 7 | 7192 | >200 | 10.5281/zenodo.10698129 |
| D-GlcA | 0.25 | 6 | 10 | 16 | 7102 | >200 | 10.5281/zenodo.10698129 |
| D-GlcA | 0.50 | 13 | 20 | 33 | 6927 | >200 | 10.5281/zenodo.10698129 |
| D-GlcA | 0.75 | 19 | 31 | 50 | 6767 | >200 | 10.5281/zenodo.10698129 |
| D-GlcA | 1.00 | 26 | 41 | 67 | 6609 | >200 | 10.5281/zenodo.10698129 |

[1] Equilibrium anomeric ratios were modeled according to Ref. S57
[2] `CHARMM36` and `prosECCo75` – 220 ns, `CHARMM36-NBFIX` – 210 ns. The last 200 ns were used for the analysis.

Table S17: **Carbohydrate systems used for the calculation of the free energy of puckering** The temperature in all simulations was 310.15 K.

| Carbohydrate | Force field | Num. of cations | Time [ns] | Zenodo link |
|---|---|---|---|---|
| GlcA | `CHARMM36` | 1 $Na^+$ | 200 | 10.5281/zenodo.10700652 |
| GlcA | `CHARMM36-NBFIX` | 1 $Na^+$ | 200 | 10.5281/zenodo.10700652 |
| GlcA | `prosECCo75` | 1 $Na^+$ | 200 | 10.5281/zenodo.10700652 |
| GalA | `CHARMM36` | 1 $Na^+$ | 200 | 10.5281/zenodo.10700652 |
| GalA | `CHARMM36-NBFIX` | 1 $Na^+$ | 200 | 10.5281/zenodo.10700652 |
| GalA | `prosECCo75` | 1 $Na^+$ | 200 | 10.5281/zenodo.10700652 |

#### S3.2 Osmotic Coefficient Calculations and Experimental Results for Acidic Saccharides

Osmotic coefficient calculations were employed to evaluate the performance of `prosECCo75` force field in modeling small organic molecules. We carried out simulations of twenty natural

L-amino acids and two monosaccharides, d-glucuronic and d-galacturonic acids. Three concentrations were simulated for amino acid solutions [0.5 M; 1 M; 2M], similar to the previous study.[S53] We also adopted the same experimental reference data. For monosaccharides, we carried out our own experimental measurements described in the Methods section. Five concentrations [0.1 M; 0.25 M; 0.5 M; 0.75 M; 1 M] were modeled to cover an experimentally measured concentration range up to 1 M. The monosaccharides were simulated in ∼2:3 ratio of $\alpha/\beta$ monomers in accord with the experimentally measured anomeric equilibrium.[S57] In the case of charged species, the system was charge-neutralized by counter ions (sodium or chloride). For computational details, see section S2.1.

#### S3.3 Experimental Determination Osmotic Coefficient for Acidic Saccharides

For saccharides, the determined experimental osmolalities were fitted to a three-parameter equation:

$$\mathrm{Osm} = 2 \cdot m_{\mathrm{X-Na}} + A \cdot (m_{\mathrm{X-Na}})^{\frac{3}{2}} + B \cdot (m_{\mathrm{X-Na}})^2 + C \cdot (m_{\mathrm{X-Na}})^3 \tag{S5}$$

where $m_{\mathrm{X-Na}}$ is molality of saccharide–sodium salt and $A$, $B$, and $C$ are adjustable parameters. Note that $2 \cdot m_{\mathrm{X-Na}}$ term is the osmolality of the ideal solution.

The osmotic coefficients were calculated from osmolality:

$$\phi = \frac{\mathrm{Osm}}{2 \cdot m_{\mathrm{X-Na}}} \tag{S6}$$

Combining the two previous equations, the osmotic coefficient can be calculated as:

$$\phi = 1 + \frac{A}{2} \cdot (m_{\mathrm{X-Na}})^{\frac{1}{2}} + \frac{B}{2} \cdot m_{\mathrm{X-Na}} + \frac{C}{2} \cdot (m_{\mathrm{X-Na}})^2 \tag{S7}$$

Parameters $A$, $B$, and $C$ used for the fitting are given in Table S18, while tabulated values of both osmolalities and osmotic coefficients are given in Table S20.

Table S18: Fitting parameters in calculating experimental osmotic coefficients of saccharide solutions.

| Saccharide | $A$ [kg$^{1/2}$/mol$^{1/2}$] | $B$ [kg/mol] | $C$ [kg$^2$/mol$^2$] |
|---|---|---|---|
| D-GalA | –0.307 | –0.508 | 0.326 |
| D-GlcA | –0.378 | 0.00822 | 0 |

#### S3.4 `prosECCo75` does not Compromise the Original CHARMM Saccharide Force Field Quality

The free energy of puckering of glucuronic acid (GlcA) and galacturonic acid (GalA) was calculated for the three studied force fields, `prosECCo75`, `CHARMM36`, and `CHARMM36-NBFIX` to evaluate the force field effects in the carbohydrate internal conformation. The composition of the systems can be found in Table S17. The free energy was calculated using well-tempered metadynamics[S58] in gromacs version 2020.4 modified with plumed.[S59,S60] The three cartesian coordinates of the saccharide pseudorotation were used as a collective variable[S61,S62] and biased by adding hills with a width of 0.01 nm and an initial height of 2kJ/mol every 250 steps. The resulting free energy was converted to polar coordinates following the methodology outlined in Ref. S63.

There are negligible structural differences between saccharides when using `CHARMM36`, `CHARMM36-NBFIX`, and `prosECCo75` force fields, see Figure S23.

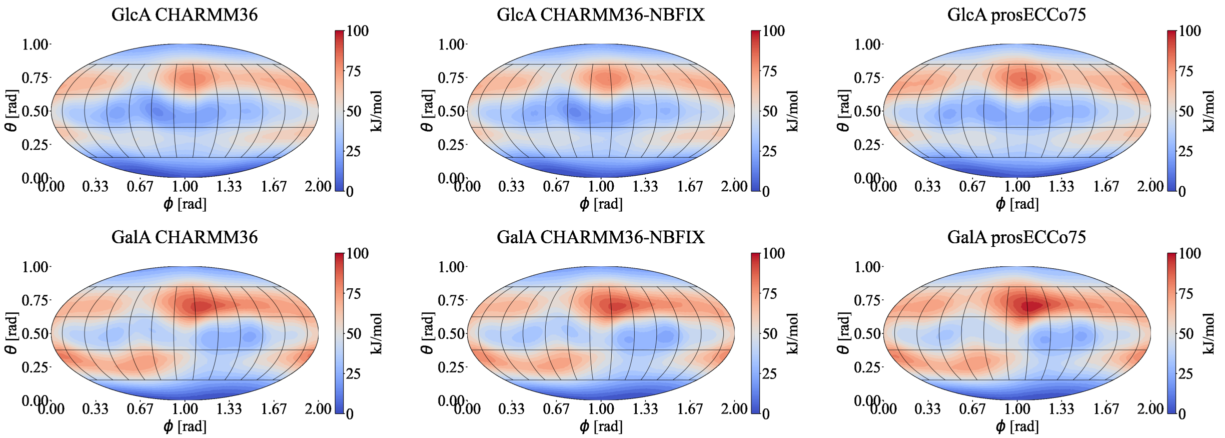


Figure S23: Puckering free energy profiles for GlcA and GalA using `CHARMM36`, `CHARMM36-NBFIX`, and `prosECCo75`.

#### S3.5 Data Osmotic Coefficient Calculation and Experiments Saccharides

Table S19: **Tabulated osmotic coefficients from simulations of saccharide solutions**

| | CHARMM36 | | CHARMM36-NBFIX | | prosECCo75 | |
|---|---|---|---|---|---|---|
| **Solution** | **Molality [m]** | **Osmotic Coefficient** | **Molality [m]** | **Osmotic Coefficient** | **Molality [m]** | **Osmotic Coefficient** |
| D-GalA–Na | 0.106 | 0.820 ± 0.026 | 0.106 | 0.935 ± 0.027 | 0.107 | 0.955 ± 0.026 |
| D-GalA–Na | 0.246 | 0.791 ± 0.028 | 0.246 | 0.967 ± 0.019 | 0.247 | 0.905 ± 0.020 |
| D-GalA–Na | 0.522 | 0.561 ± 0.030 | 0.520 | 0.915 ± 0.014 | 0.524 | 0.863 ± 0.016 |
| D-GalA–Na | 0.814 | 0.440 ± 0.026 | 0.807 | 0.893 ± 0.013 | 0.817 | 0.806 ± 0.011 |
| D-GalA–Na | 1.125 | 0.361 ± 0.010 | 1.111 | 0.870 ± 0.010 | 1.130 | 0.732 ± 0.013 |
| D-GlcA–Na | 0.106 | 0.899 ± 0.033 | 0.106 | 0.952 ± 0.029 | 0.107 | 0.932 ± 0.027 |
| D-GlcA–Na | 0.246 | 0.845 ± 0.019 | 0.246 | 0.962 ± 0.022 | 0.247 | 0.949 ± 0.018 |
| D-GlcA–Na | 0.521 | 0.737 ± 0.014 | 0.520 | 0.919 ± 0.016 | 0.524 | 0.883 ± 0.015 |
| D-GlcA–Na | 0.813 | 0.531 ± 0.016 | 0.807 | 0.916 ± 0.014 | 0.818 | 0.861 ± 0.010 |
| D-GlcA–Na | 1.123 | 0.447 ± 0.014 | 1.110 | 0.918 ± 0.012 | 1.130 | 0.829 ± 0.010 |

Table S20: **Experimental measurements of osmotic coefficients of saccharide solutions** The temperature in all experiments was 310.15 K.

| Solution | Molality [m] | Osmolality [m] | Osmotic Coefficient | Solution | Molality [m] | Osmolality [m] | Osmotic Coefficient |
|---|---|---|---|---|---|---|---|
| D-GalA–Na | 0.0495 | 0.095 ± 0.004 | 0.9626 ± 0.041 | D-GlcA–Na | 0.0496 | 0.096 ± 0.006 | 0.9668 ± 0.061 |
| D-GalA–Na | 0.0996 | 0.189 ± 0.002 | 0.9484 ± 0.011 | D-GlcA–Na | 0.0761 | 0.143 ± 0.007 | 0.9401 ± 0.047 |
| D-GalA–Na | 0.1243 | 0.229 ± 0.004 | 0.9222 ± 0.017 | D-GlcA–Na | 0.0917 | 0.170 ± 0.005 | 0.9269 ± 0.030 |
| D-GalA–Na | 0.1537 | 0.279 ± 0.011 | 0.9076 ± 0.036 | D-GlcA–Na | 0.1460 | 0.274 ± 0.009 | 0.9386 ± 0.031 |
| D-GalA–Na | 0.2003 | 0.360 ± 0.003 | 0.8985 ± 0.008 | D-GlcA–Na | 0.1982 | 0.365 ± 0.005 | 0.9205 ± 0.013 |
| D-GalA–Na | 0.2192 | 0.383 ± 0.005 | 0.8738 ± 0.011 | D-GlcA–Na | 0.2494 | 0.457 ± 0.006 | 0.9163 ± 0.013 |
| D-GalA–Na | 0.2472 | 0.428 ± 0.007 | 0.8659 ± 0.015 | D-GlcA–Na | 0.2991 | 0.537 ± 0.011 | 0.8976 ± 0.019 |
| D-GalA–Na | 0.2883 | 0.490 ± 0.004 | 0.8496 ± 0.007 | D-GlcA–Na | 0.3884 | 0.692 ± 0.009 | 0.8909 ± 0.012 |
| D-GalA–Na | 0.3982 | 0.661 ± 0.010 | 0.8300 ± 0.013 | D-GlcA–Na | 0.4974 | 0.865 ± 0.011 | 0.8696 ± 0.012 |
| D-GalA–Na | 0.4986 | 0.809 ± 0.010 | 0.8113 ± 0.011 | D-GlcA–Na | 0.5890 | 1.002 ± 0.015 | 0.8506 ± 0.013 |
| D-GalA–Na | 0.5754 | 0.903 ± 0.012 | 0.7847 ± 0.011 | D-GlcA–Na | 0.7372 | 1.237 ± 0.023 | 0.8390 ± 0.016 |
| D-GalA–Na | 0.7680 | 1.182 ± 0.011 | 0.7696 ± 0.007 | D-GlcA–Na | 0.9104 | 1.503 ± 0.021 | 0.8255 ± 0.012 |
| D-GalA–Na | 0.8805 | 1.334 ± 0.012 | 0.7575 ± 0.007 | | | | |

### S4 Supplementary Information Ions

#### S4.1 Partial Charges for Ions

`prosECCo75` force fields development includes 3 new ions to cover the two missing halides, $Br^-$ (2 variants: Br_s and Br_2s), and $I^-$ (I_s). Both sets we designed to align reasonably well with the other ions in the series, see Figure S24.

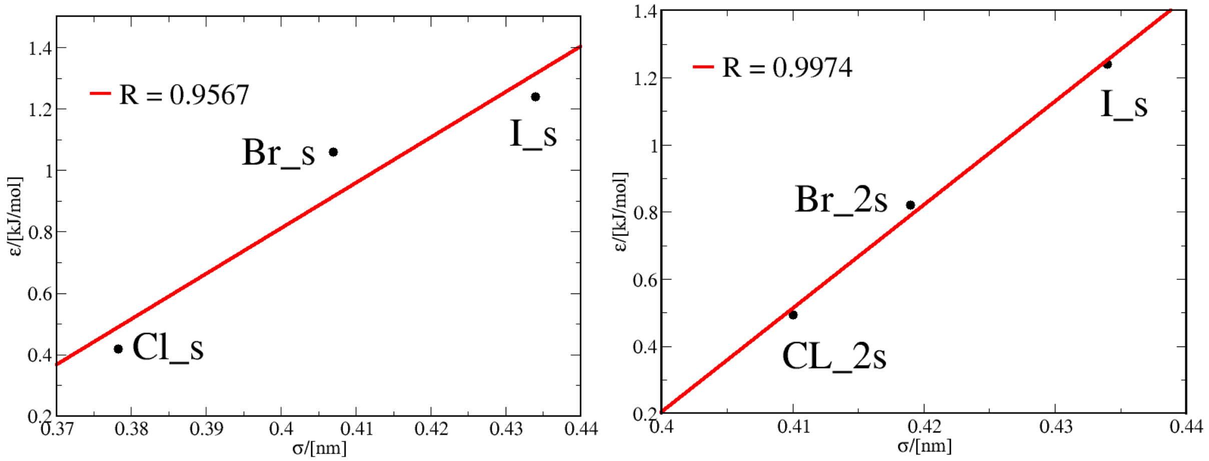


(a) Anions”_s”

(b) Anions”_2s”

Figure S24: Relation between $\alpha$ and $\epsilon$ parameters for ions in `prosECCo75`: (a) monovalent halogens”_s” and (b) monovalent halogens”_2s”. $I^-$ has only a variant compatible with both ”_s” and _2s” variants.

For the parameter optimization process, we use SPC/E [S64] water since the default TIP3P[S65] model has a wrong density at biological conditions and cannot be compared with experiments.

#### S4.2 Simulated Ions

The molecular details of the simulated ion systems can be found in Table S21. The ion optimization was done using the Noosé-Hoover thermostat and the Parrinello-Rahman barostat with relaxation times of 1.0 ps and 3.0 ps, respectively. Temperature was fixed at 298 K and pressure to 1 bar. Additionally, the barostat used a compressibility of 5E-5 bar$^{-1}$. The cut-off schemes used were PME schemes (both Coulomb and Lennard-Jones) to take into account non-bonding interactions after the cut-off radius of 1.2 nm. The simulation time was 21 ns, where the first nanosecond was used as equilibration and a time step of 2 fs. The sampling frequency of saving snapshots was 10 ps, analyzing a total of 2000 configurations per hydration shell and density value.

Table S21: **Salt systems used for the optimization of the monovalent salts ($K^+X^-$), where $X^-$ may be $Cl^-$, $Br^-$, $I^-$.**

| Salt | Number of salt molecules | Number of water molecules | Concentration [m] | Time [ns] | Temp [K] |
|---|---|---|---|---|---|
| $K^+X^-$ | 25 | 2776 | 0.5 | 21 | 298. |
| $K^+X^-$ | 100 | 2776 | 2.0 | 21 | 298. |
| $K^+X^-$ | 200 | 2776 | 4.0 | 21 | 298. |

#### S4.3 Comparison Neutron Scattering Data and Simulation for Ion Solutions

In water solutions, the main contributions of the scattering pattern are the hydrogen and oxygen atoms of the water molecule. The hydrogen contribution can be removed if the experiments take place in null water, which is water with a ratio of 1.784:1 of light and heavy water ($H_2O$:$D_2O$) because the nuclei have an opposite contribution to the neutron scattering experiment. For the liquids measured in this study, the constitution of the total scattering patterns ($S$) is:

$$\begin{aligned}
{}^{\text{water}}S(Q) &= 37.4S_{OO}^{pure}(Q) - 37.4 \\
{}^{\text{KCl}}S(Q) &= 34.1S_{OO}(Q) + 8.1S_{OCl}(Q) + 3.1S_{OK}(Q)+ \\
&\quad + 0.5S_{ClCl}(Q) + 0.4S_{KCl}(Q) + 0.1S_{KK}(Q) - 46.3 \\
{}^{\text{KBr}}S(Q) &= 34.1S_{OO}(Q) + 5.7S_{OBr}(Q) + 3.1S_{OK}(Q)+ \\
&\quad + 0.2S_{BrBr}(Q) + 0.3S_{KBr}(Q) + 0.1S_{KK}(Q) - 43.6 \\
{}^{\text{KI}}S(Q) &= 34.1S_{OO}(Q) + 4.4S_{OI}(Q) + 3.1S_{OK}(Q)+ \\
&\quad + 0.2S_{II}(Q) + 0.2S_{KI}(Q) + 0.1S_{KK}(Q) - 42.1
\end{aligned} \tag{S8}$$

where the prefactors (in millibarns) are calculated from products of the atom concentrations of the two nuclei in question and the coherent neutron scattering length of the two nuclei in question, see explanation in Reference S66. These may be Fourier transformed to yield radial distribution functions ($G$)that can be directly compared to molecular dynamics simulations:

$$\begin{aligned}
{}^{\text{water}}G(Q) &= 37.5G_{OO}^{pure}(Q) - 37.5 \\
{}^{\text{KCl}}G(Q) &= 34.1G_{OO}(Q) + 8.1G_{OCl}(Q) + 3.1G_{OK}(Q)+ \\
&\quad + 0.5G_{ClCl}(Q) + 0.4G_{KCl}(Q) + 0.1G_{KK}(Q) - 46.3 \\
{}^{\text{KBr}}G(Q) &= 34.1G_{OO}(Q) + 5.7G_{OBr}(Q) + 3.1G_{OK}(Q)+ \\
&\quad + 0.2G_{BrBr}(Q) + 0.3G_{KBr}(Q) + 0.1G_{KK}(Q) - 43.6 \\
{}^{\text{KI}}G(Q) &= 34.1G_{OO}(Q) + 4.4G_{OI}(Q) + 3.1G_{OK}(Q)+ \\
&\quad + 0.2G_{II}(Q) + 0.2G_{KI}(Q) + 0.1G_{KK}(Q) - 42.1
\end{aligned} \tag{S9}$$

Noting there are some differences between $S_{OO}^{pure}(Q)$ for pure water and the halide solutions, subtracting 0.910 $^{water}G(r)$ from the structure factor of the halide solution largely cancels the Oxygen-Oxygen (OO) term and highlights the remaining structure of the solution, which is principally constituted of the O-Halide, and O-K terms with a smaller contribution from the ion-ion terms. This exact function can be calculated from MD simulations and compared with the experiment. In each case the function compared is $^{KCl}\Delta G = {}^{KCl}G(r) - 0.910 \cdot^{water} G(r)$, $^{KBr}\Delta G = {}^{KBr}G(r) - 0.910 \cdot {}^{water}G(r)$ and $^{KI}\Delta G = {}^{KI}G(r) - 0.910 \cdot {}^{water}G(r)$.

To compare the simulation details with the experimental neutron scattering parameters, the individual simulation pair RDFs were scaled using the prefactors provided in Table S9.

#### S4.4 Characterization "_s" Anion Series and K_s Cation

The parameterization of the $Br^-$ and $I^-$ anions was done to improve the density against the $K^+$ (K_s). To this end, the SPC/E water model[S64] was used as it provides better pure density than the CHARMM36 default mTIP3P water model.[S65] Radial distributions for The _s anion series can be found in Figure S25. The performance of the "_s" and "_2s" models with respect to the experimental density can be found in Table S22.

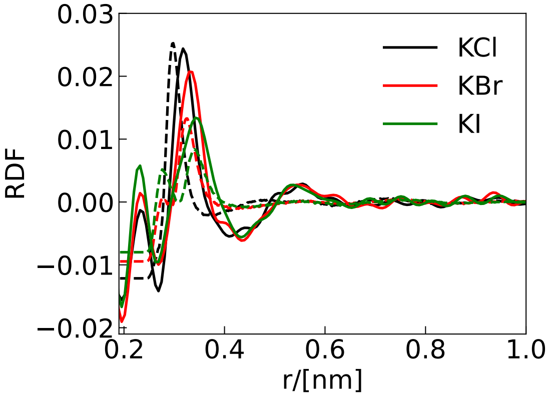


(a) Anions"_s" solvation shell.

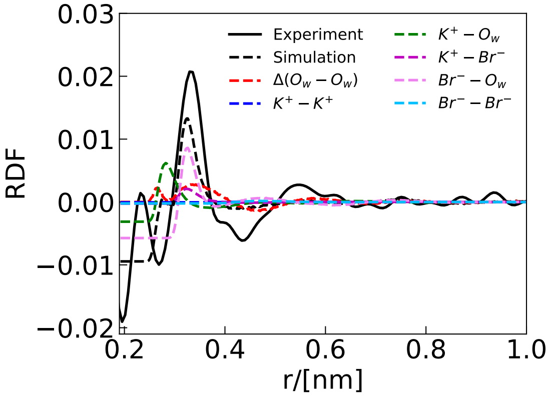


(b) Contributions for Br_s with K_s solution.

Figure S25: Characterization solvation shell "_s" series. a) Solvation shell of 4m potassium chloride (KCl), 4m potassium bromide (KBr), and 4m potassium iodide (KI). The solid lines are the experimental neutron scattering results, and the dashed lines are the simulation results using K_s, Cl_s, Br_s, and I_s parameters. Right: Scaled contribution by the prefactor for each atomic pair (Equation S9) and shifted to zero in Br_s–K_s solution.

Table S22: Density values for KCl, KBr, and KI using the K_s, Cl_s, Cl_2s, Br_s, Br_2s, and I_s models with SPC/E at different concentrations at 298 K and 1 bar. Experimental values[S67] and their differences are provided.

| | | Density/[ kg/m$^3$] | |
|---|---|---|---|
| conc/[M] | experimental | Simulation | Difference |
| K_s Cl_s | | | |
| 0.5 | 1019.73 | 1019.03 | -0.70 |
| 2.0 | 1081.77 | 1072.90 | -8.87 |
| 4.0 | 1152.25 | 1135.16 | -17.09 |
| K_s Cl_2s | | | |
| 0.5 | 1019.73 | 1013.57 | -6.16 |
| 2.0 | 1081.77 | 1052.76 | -29.01 |
| 4.0 | 1152.25 | 1098.24 | -54.01 |
| K_s Br_s | | | |
| 0.5 | 1038.16 | 1034.30 | -3.86 |
| 2.0 | 1151.08 | 1130.20 | -20.88 |
| 4.0 | 1280.94 | 1241.56 | -39.38 |
| K_s Br_2s | | | |
| 0.5 | 1038.156 | 1032.47 | -5.69 |
| 2.0 | 1151.081 | 1124.42 | -26.66 |
| 4.0 | 1280.935 | 1230.16 | -50.78 |
| K_s I_s | | | |
| 0.5 | 1055.50 | 1052.66 | -2.84 |
| 2.0 | 1213.56 | 1197.38 | -16.18 |
| 4.0 | 1391.38 | 1361.84 | -29.54 |

### References


(S1) Nencini, R. Development of PROSECCO PC membranes - PN model, POPC membranes, different calcium and sodium concentrations, 310K. 2019.

(S2) Nencini, R. POPC bilayer at 310K with the prosECCo75 force field, 111 mM NaCl . 2024; https://doi.org/10.5281/zenodo.10464960.

(S3) Nencini, R. CHARMM36, NB-Fix approaches, without NBFIX, POPC membrane, Ca, Na ions,. 2019.

(S4) Nencini, R. POPC bilayer at 310K with the CHARMM36-NBFIX force field, 111 mM CaCl2. 2024; https://doi.org/10.5281/zenodo.10464946.

(S5) Nencini, R. POPC bilayer at 310K with the CHARMM36-NBFIX force field, 111 mM NaCl . 2024; https://doi.org/10.5281/zenodo.10464958.

(S6) Nencini, R. POPC bilayer at 310K with the CHARMM36 force field, 111 mM CaCl2 . 2024; https://doi.org/10.5281/zenodo.10464498.

(S7) Nencini, R. POPC bilayer at 310K with the CHARMM36 force field, 111 mM NaCl . 2024; https://doi.org/10.5281/zenodo.10464448.

(S8) Nencini, R. Development of PROSECCO PC membranes - PN model, melting temperature, DPPC membranes. 2019; https://doi.org/10.5281/zenodo.3332640.

(S9) Nencini, R. MD Simulation of Calcium binding to DPPC bilayers, prosECCo75 FF, 450mM. 2019; https://doi.org/10.5281/zenodo.3371326.

(S10) Martinez-Seara, H. prosECCo75 paper related MD Simulation, DPPC prosECCo75 LO, melting T . 2024; https://doi.org/10.5281/zenodo.10635500.

(S11) Martinez-Seara, H. prosECCo75 paper related MD Simulation, DPPC prosECCo75 GEL, melting T . 2024; https://doi.org/10.5281/zenodo.10635502.

(S12) Martinez-Seara, H. prosECCo75 paper related MD Simulation, DPPC CHARMM LO, melting T . 2024; https://doi.org/10.5281/zenodo.10635496.

(S13) Martinez-Seara, H. prosECCo75 paper related MD Simulation, DPPC CHARMM GEL, melting T . 2024; https://doi.org/10.5281/zenodo.10635498.

(S14) Nencini, R. MD Simulation of POPS 144 144 Na contraions prosECCo75. 2024; https://doi.org/10.5281/zenodo.10467181.

(S15) Javanainen, M. MD Simulation of Calcium binding to POPC:POPS 5:1 bilayers, prosECCo75 FF, K+ contraions. 2024; https://doi.org/10.5281/zenodo.10683998.

(S16) Nencini, R. MD Simulation of POPC 120 POPS 24 Na contraions prosECCo75. 2024; https://doi.org/10.5281/zenodo.10467179.

(S17) Nencini, R. MD Simulation of POPC 120 POPS 24 Na contraions 100mM CaCl prosECCo75. 2024; https://doi.org/10.5281/zenodo.10467165.

(S18) Nencini, R. MD Simulation of POPC 120 POPS 24 Na contraions 300mM CaCl prosECCo75. 2024; https://doi.org/10.5281/zenodo.10467173.

(S19) Nencini, R. MD Simulation of POPC 120 POPS 24 Na contraions 1000mM CaCl prosECCo75. 2024; https://doi.org/10.5281/zenodo.10467161.

(S20) Nencini, R. MD Simulation of POPC 120 POPS 24 Na contraions 3000mM CaCl prosECCo75. 2024; https://doi.org/10.5281/zenodo.10467171.

(S21) Antila, H.; Buslaev, P.; Favela-Rosales, F.; Ferreira, T. M.; Gushchin, I.; Javanainen, M.; Kav, B.; Madsen, J. J.; Melcr, J.; Miettinen, M. S.; Määttä, J.; Nencini, R.; Ollila, O. H. S.; Piggot, T. J. Headgroup Structure and Cation Binding in Phosphatidylserine Lipid Bilayers. *J. Phys. Chem. B* **2019**, *123*, 9066–9079.

(S22) Nencini, R. POPC 120 POPS 24 Na counterions CHARMM NBFIX REMOVED 298K. 2024; https://doi.org/10.5281/zenodo.10700845.

(S23) Nencini, R. POPC 120 POPS 24 Na counterions 100mM CaCl CHARMM NBFIX REMOVED 298K. 2024; https://doi.org/10.5281/zenodo.10700796.

(S24) Nencini, R. POPC 120 POPS 24 Na counterions 300mM CaCl CHARMM NBFIX REMOVED 298K. 2024; https://doi.org/10.5281/zenodo.10700778.

(S25) Nencini, R. POPC 120 POPS 24 Na counterions 1000mM CaCl CHARMM NBFIX REMOVED 298K. 2024; https://doi.org/10.5281/zenodo.10700782.

(S26) Nencini, R. POPC 120 POPS 24 Na counterions 3000mM CaCl CHARMM NBFIX REMOVED 298K. 2024; https://doi.org/10.5281/zenodo.10700786.

(S27) Nencini, R. PC/PS 5:1 bilayer, Na contraions, additional CaCl_2, CHARMM36 FF, NBFIX-Han. 2019; https://doi.org/10.5281/zenodo.3371326.

(S28) Nencini, R. POPC 120 POPS 24 Na counterions CHARMM NBFIX HAN 298K. 2024; https://doi.org/10.5281/zenodo.10730144.

(S29) Nencini, R. POPC 120 POPS 24 Na counterions 100mM CaCl CHARMM NBFIX HAN 298K. 2024; https://doi.org/10.5281/zenodo.10730166.

(S30) Nencini, R. POPC 120 POPS 24 Na counterions 300mM CaCl CHARMM NBFIX HAN 298K. 2024; https://doi.org/10.5281/zenodo.10730150.

(S31) Nencini, R. POPC 120 POPS 24 Na counterions 1000mM CaCl CHARMM NBFIX HAN 298K. 2024; https://doi.org/10.5281/zenodo.10730157.

(S32) Nencini, R. POPC 120 POPS 24 Na counterions 3000mM CaCl CHARMM NBFIX HAN 298K. 2024; https://doi.org/10.5281/zenodo.10730163.

(S33) Javanainen, M. MD Simulation of POPC cholesterol bilayers, prosECCo75 FF, different compositions. 2024; https://doi.org/10.5281/zenodo.10683741.

(S34) Ferreira, T. M.; Coreta-Gomes, F.; Ollila, O. S.; Moreno, M. J.; Vaz, W. L.; Topgaard, D. Cholesterol and POPC Segmental Order Parameters in Lipid Membranes: Solid State $^{1}$H–$^{13}$C NMR and MD Simulation Studies. *Phys. Chem. Chem. Phys.* **2013**, *15*, 1976–1989.

(S35) Bacle, A.; Buslaev, P.; Garcia-Fandino, R.; Favela-Rosales, F.; Mendes Ferreira, T.; Fuchs, P. F.; Gushchin, I.; Javanainen, M.; Kiirikki, A. M.; Madsen, J. J.; Melcr, J.; Rodriguez, P. M.; Miettinen, M. S.; Ollila, O. H. S.; Papadopoulos, C. G.; Peón, A.;

Piggot, T. J.; Piñeiro, A.; Virtanen, S. I. Inverse Conformational Selection in Lipid–Protein Binding. *J. Am. Chem. Soc.* **2021**, *143*, 13701–13709.

(S36) Melcr, J.; Ferreira, T. M.; Jungwirth, P.; Ollila, O. S. Improved Cation Binding to Lipid Bilayers With Negatively Charged POPS by Effective Inclusion of Electronic Polarization. *J. Chem. Theory Comput.* **2019**, *16*, 738–748.

(S37) Klauda, J. B.; Venable, R. M.; Freites, J. A.; O'Connor, J. W.; Tobias, D. J.; Mondragon-Ramirez, C.; Vorobyov, I.; MacKerell Jr, A. D.; Pastor, R. W. Update of the CHARMM All-Atom Additive Force Field for Lipids: Validation on Six Lipid Types. *J. Phys. Chem. B* **2010**, *114*, 7830–7843.

(S38) Kučerka, N.; Nieh, M.-P.; Katsaras, J. Fluid Phase Lipid Areas and Bilayer Thicknesses of Commonly Used Phosphatidylcholines as a Function of Temperature. *Biochim. Biophys. Acta* **2011**, *1808*, 2761–2771.

(S39) Pan, J.; Cheng, X.; Monticelli, L.; Heberle, F. A.; Kučerka, N.; Tieleman, D. P.; Katsaras, J. The Molecular Structure of a Phosphatidylserine Bilayer Determined by Scattering and Molecular Dynamics Simulations. *Soft Matter* **2014**, *10*, 3716–3725.

(S40) Kučerka, N.; van Oosten, B.; Pan, J.; Heberle, F. A.; Harroun, T. A.; Katsaras, J. Molecular Structures of Fluid Phosphatidylethanolamine Bilayers Obtained From Simulation-To-Experiment Comparisons and Experimental Scattering Density Profiles. *J. Phys. Chem. B* **2015**, *119*, 1947–1956.

(S41) Kučerka, N.; Tristram-Nagle, S.; Nagle, J. F. Structure of Fully Hydrated Fluid Phase Lipid Bilayers With Monounsaturated Chains. *J. Membr. Biol.* **2006**, *208*, 193–202.

(S42) Kučerka, N.; Nagle, J. F.; Sachs, J. N.; Feller, S. E.; Pencer, J.; Jackson, A.; Katsaras, J. Lipid Bilayer Structure Determined by the Simultaneous Analysis of Neutron and X-Ray Scattering Data. *Biophys. J.* **2008**, *95*, 2356–2367.

(S43) Javanainen, M. Simulation of a POPE bilayer at 310K with the CHARMM36 force field. 2019; `https://doi.org/10.5281/zenodo.2641987`.

(S44) Tempra, C.; Ollila, O. H. S.; Javanainen, M. Accurate Simulations of Lipid Monolayers Require a Water Model With Correct Surface Tension. *J. Chem. Theory Comput.* **2022**, *18*, 1862–1869.

(S45) Javanainen, M.; Lamberg, A.; Cwiklik, L.; Vattulainen, I.; Ollila, O. S. Atomistic Model for Nearly Quantitative Simulations of Langmuir Monolayers. *Langmuir* **2018**, *34*, 2565–2572.

(S46) Van Meer, G.; Voelker, D. R.; Feigenson, G. W. Membrane Lipids: Where They Are and How They Behave. *Nat. Rev. Mol. Cell Biol.* **2008**, *9*, 112.

(S47) Brown, M. F.; Seelig, J. Influence of Cholesterol on the Polar Region of Phosphatidylcholine and Phosphatidylethanolamine Bilayers. *Biochemistry* **1978**, *17*, 381–384.

(S48) Botan, A.; Favela-Rosales, F.; Fuchs, P. F.; Javanainen, M.; Kanduč, M.; Kulig, W.; Lamberg, A.; Loison, C.; Lyubartsev, A.; Miettinen, M. S.; Monticelli, L.; Määttä, J.; Ollila, O. H. S.; Retegan, M.; Rog, T.; Santuz, H.; Tynkkynen, J. Toward Atomistic Resolution Structure of Phosphatidylcholine Headgroup and Glycerol Backbone at Different Ambient Conditions. *J. Phys. Chem. B* **2015**, *119*, 15075–15088.

(S49) Javanainen, M.; Melcrová, A.; Magarkar, A.; Jurkiewicz, P.; Hof, M.; Jungwirth, P.; Martinez-Seara, H. Two Cations, Two Mechanisms: Interactions of Sodium and Calcium With Zwitterionic Lipid Membranes. *Chem. Commun.* **2017**, *53*, 5380–5383.

(S50) Altenbach, C.; Seelig, J. Calcium Binding to Phosphatidylcholine Bilayers as Studied by Deuterium Magnetic Resonance. Evidence for the Formation of a Calcium Complex With Two Phospholipid Molecules. *Biochemistry* **1984**, *23*, 3913–3920.

(S51) Melcr, J.; Martinez-Seara, H.; Nencini, R.; Kolafa, J.; Jungwirth, P.; Ollila, O. S. Accurate Binding of Sodium and Calcium to a POPC Bilayer by Effective Inclusion of Electronic Polarization. *J. Phys. Chem. B* **2018**, *122*, 4546–4557.

(S52) Akutsu, H.; Seelig, J. Interaction of Metal Ions With Phosphatidylcholine Bilayer Membranes. *Biochemistry* **1981**, *20*, 7366–7373.

(S53) Miller, M. S.; Lay, W. K.; Elcock, A. H. Osmotic Pressure Simulations of Amino Acids and Peptides Highlight Potential Routes to Protein Force Field Parameterization. *J. Phys. Chem. B* **2016**, *120*, 8217–8229.

(S54) Luo, Y.; Roux, B. Simulation of Osmotic Pressure in Concentrated Aqueous Salt Solutions. *J. Phys. Chem. Lett.* **2009**, *1*, 183–189.

(S55) Lay, W. K.; Miller, M. S.; Elcock, A. H. Reparameterization of Solute—Solute Interactions for Amino Acid–Sugar Systems Using Isopiestic Osmotic Pressure Molecular Dynamics Simulations. *J. Chem. Theory Comput.* **2017**, *13*, 1874–1882.

(S56) Miller, M. S.; Lay, W. K.; Li, S.; Hacker, W. C.; An, J.; Ren, J.; Elcock, A. H. Reparametrization of Protein Force Field Nonbonded Interactions Guided by Osmotic Coefficient Measurements From Molecular Dynamics Simulations. *J. Chem. Theory Comput.* **2017**, *13*, 1812–1826.

(S57) Kerins, L.; Byrne, S.; Gabba, A.; Murphy, P. V. Anomer Preferences for Glucuronic and Galacturonic Acid and Derivatives and Influence of Electron-Withdrawing Substituents. *J. Org. Chem.* **2018**, *83*, 7714–7729.

(S58) Barducci, A.; Bussi, G.; Parrinello, M. Well-Tempered Metadynamics: A Smoothly Converging and Tunable Free-Energy Method. *Phys. Rev. Lett.* **2008**, *100*.

(S59) Abraham, M. J.; Murtola, T.; Schulz, R.; Páll, S.; Smith, J. C.; Hess, B.; Lindahl, E.

GROMACS: High Performance Molecular Simulations Through Multi-Level Parallelism From Laptops to Supercomputers. *SoftwareX* **2015**, *1-2*, 19–25.

(S60) Tribello, G. A.; Bonomi, M.; Branduardi, D.; Camilloni, C.; Bussi, G. Plumed 2: New Feathers for an Old Bird. *Comput. Phys. Commun.* **2014**, *185*, 604–613.

(S61) Cremer, D.; Pople, J. A. A General Definition of Ring Puckering Coordinates. *J. Am. Chem. Soc.* **1975**, *97*, 1354–1358.

(S62) Biarnés, X.; Ardèvol, A.; Planas, A.; Rovira, C.; Laio, A.; Parrinello, M. The Conformational Free Energy Landscape of $\beta$-D-Glucopyranose. Implications for Substrate Preactivation in $\beta$-Glucoside Hydrolases. *J. Am. Chem. Soc.* **2007**, *129*, 10686–10693.

(S63) Palivec, V.; Johannessen, C.; Kaminský, J.; Martinez-Seara, H. Use of Raman and Raman Optical Activity to Extract Atomistic Details of Saccharides in Aqueous Solution. *PLoS Comput. Biol.* **2022**, *18*, e1009678.

(S64) Chatterjee, S.; Debenedetti, P. G.; Stillinger, F. H.; Lynden-Bell, R. M. A Computational Investigation of Thermodynamics, Structure, Dynamics and Solvation Behavior in Modified Water Models. *J. Chem. Phys.* **2008**, *128*.

(S65) MacKerell, A. D.; Bashford, D.; Bellott, M.; Dunbrack, R. L.; Evanseck, J. D.; Field, M. J.; Fischer, S.; Gao, J.; Guo, H.; Ha, S.; Joseph-McCarthy, D.; Kuchnir, L.; Kuczera, K.; Lau, F. T. K.; Mattos, C.; Michnick, S.; Ngo, T.; Nguyen, D. T.; Prodhom, B.; Reiher, W. E.; Roux, B.; Schlenkrich, M.; Smith, J. C.; Stote, R.; Straub, J.; Watanabe, M.; Wiórkiewicz-Kuczera, J.; Yin, D.; Karplus, M. All-Atom Empirical Potential for Molecular Modeling and Dynamics Studies of Proteins. *J. Phys. Chem. B* **1998**, *102*, 3586–3616.

(S66) Mason, P. E.; Ansell, S.; Neilson, G. W. Neutron Diffraction Studies of Electrolytes in Null Water: A Direct Determination of the First Hydration Zone of Ions. *J. Phys. Condens. Matter* **2006**, *18*, 8437–8447.

(S67) Laliberté, M. A Model for Calculating the Heat Capacity of Aqueous Solutions, With Updated Density and Viscosity Data. *J. Chem. Eng. Data* **2009**, *54*, 1725–1760.
